# A central role of glutamine in *Chlamydia* infection

**DOI:** 10.1101/742817

**Authors:** Karthika Rajeeve, Nadine Vollmuth, Sudha Janaki-Raman, Thomas Wulff, Maximilian Schmalhofer, Werner Schmitz, Apoorva Baluapuri, Claudia Huber, Julian Fink, Francesca R. Dejure, Elmar Wolf, Wolfgang Eisenreich, Almut Schulze, Jürgen Seibel, Thomas Rudel

## Abstract

Obligate intracellular bacteria like *Chlamydia trachomatis* undergo a complex developmental cycle between infectious non-replicative (EBs) and non-infectious replicative (RBs) forms. EBs shortly after entering a host cell transform to RBs, a crucial process in infection, initiating chlamydial replication. As *Chlamydia* fail to replicate outside the host cell it is currently unknown how the transition from EBs to RBs is initiated. Here we show in a cell-free approach in axenic media that uptake of glutamine by the bacteria is critical to initiate EB-RB transition. These bacteria utilize glutamine to synthesize cell wall peptidoglycan which has recently been detected in the septa of replicating intracellular *Chlamydia.* The increased requirement for glutamine in infected cells is achieved by reprogramming the glutamine metabolism in a c-Myc-dependent manner. Glutamine was effectively taken up by the glutamine transporter SLC1A5 and metabolized via glutaminase. Interference with this metabolic reprogramming limited growth of *Chlamydia.* Intriguingly, *Chlamydia* failed to produce progeny in SLC1A5 knockout mice. Thus, we report on the central role of glutamine for the development of an obligate intracellular pathogenic bacterium and the reprogramming of host glutamine metabolism, which may provide a basis for innovative anti-infective strategies.

---

*Chlamydia trachomatis* is an obligate intracellular bacterium and with more than 130 million infections the most frequent cause of bacterial sexually transmitted diseases ^1^. In a biphasic developmental cycle, EBs enter the host cell and convert then into RBs, which undergo replication via binary fission in a membrane-enclosed structure called inclusion. Later during the infection cycle, RBs convert back into EBs, which on lysis of the cell can readily infect neighboring cells. For a long time, it was thought that *Chlamydia* do not form peptidoglycan until peptidoglycan ring-like structures were recently discovered exclusively in the septa of dividing RBs ^2, 3^.

Due to its obligate intracellular lifestyle and an evolutionary reduced genome of only ∼1 Mb ^4^, *Chlamydia* requires multiple metabolites from the host cell. Nevertheless, this pathogen encodes the enzymes required to utilize D-glucose-6-phosphate (G6P) and to convert it into pyruvate via glycolysis ^5^. Genomic data reveal that *Chlamydia* lacks the enzyme hexokinase and substrate specific components of a phosphotransferase system. Thus, the bacteria depend on G6P from the host cell. In EBs, ATP is generated from G6P either via glycolysis or oxidative phosphorylation, whereas in RBs ATP is provided by the host cell, while G6P is nearly exclusively used for cell wall biosynthesis ^6, 7^. The TCA cycle of *Chlamydia* is incomplete due to the absence of three key enzymes, citrate synthase (GltA), aconitase (Acn) and isocitrate dehydrogenase (Icd). This incomplete metabolic pathway has to be complemented by the constant metabolic supply from the host ^4^. We have previously shown that *C. trachomatis* takes up host-derived malate to feed the partial TCA cycle ^6^. In addition, amino acids have to be acquired from the host cells due to the limited capacity of *Chlamydia* to synthesize amino acids ^6^. Despite the advancements in the knowledge about chlamydial metabolism, the trigger initiating the transition of EBs to RBs after entry into cells is still unknown.

## Results

### Glutamine metabolism in *Chlamydia:* A signal for EB to RB transition

In our attempts to further define the metabolic interaction of *Chlamydia* and host cell, we performed uptake assays in axenic culture to test whether glutamine (Gln), glutamate (Glu) and α-ketoglutarate (α-KG) previously predicted to feed the partial TCA cycle of *Chlamydia* ^4, 6, 8^ are directly taken up by these bacteria (Fig. 1a,b). Whereas the amount of Gln decreased rapidly, neither Glu nor α-KG were significantly consumed, as shown by GC-MS analyses of the supernatant (Fig. 1c). Using isotopologue profiling with Gln containing only ^13^C atoms instead of ^12^C atoms ([U-^13^C_5_]Gln) as a supplement to the axenic culture, GC-MS analyses of chlamydial extracts revealed that the ^13^C label is efficiently transferred to Glu, Asp and intermediates of the partial TCA cycle, with the ^13^C-labelled fractions ranging from 80% in Gln to less than 5% in pyruvate (Fig. 1d). Substantial fractions of ^13^C-labelling in Glu (60%), Asp (50%), succinate (20%), fumarate (35%), and malate (8%) (Fig. 1d) further indicated that exogenous Gln is metabolized via glutaminolysis and TCA cycle to generate Glu, α-KG, succinate, fumarate, malate, oxaloacetate and aspartate. In line with this was the finding that the amount of chlamydial NADH probably formed by the conversion of α-KG into succinate and malate into oxaloacetate within the truncated chlamydial TCA cycle (Fig. 1a) is significantly increased when exogenous Gln is added to the axenic medium (Supplementary Fig. 1a, 1d).

**Fig 1:**
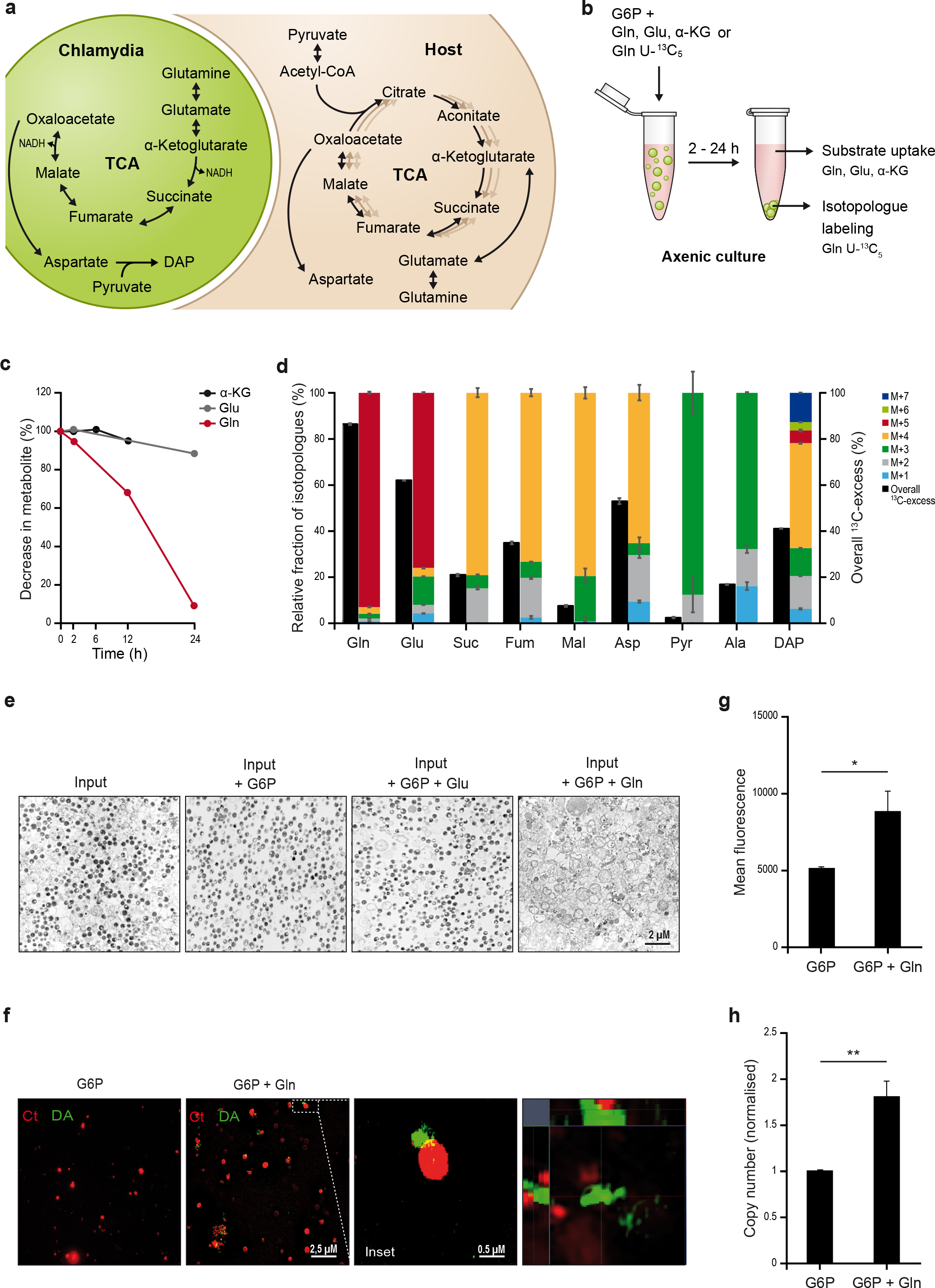
Cell wall synthesis of *Chlamydia* grown in axenic medium is induced by glutamine. **a.** Central metabolic pathways of the chlamydial and host TCA cycle. **b.** Diagram showing the setup of the experiment. **c.** The relative amounts of the indicated metabolites were determined in the supernatant of axenic cultures with *Chlamydia* at different time points by GC/MS using internal standards (see Methods). **d.** ^13^C-enrichments and isotopologue distributions of selected metabolites in *Chlamydia* cultured in axenic medium with [U-^13^C_5_]Gln for 24 h. All metabolites, with the exception of DAP, were analyzed after mechanical cell disruption in MeOH; DAP was analyzed after acid hydrolysis of the cell debris. **e.** Chlamydial EBs were purified using renografin gradient (Input) and incubated in axenic culture containing either only G6P or G6P together with Gln or Glu. After 2 h of incubation, the bacteria were pelleted and analyzed by transmission electron microscopy. Representative images from three independent experiments are shown. **f.** Purified chlamydial EBs were treated with only G6P or G6P together with Gln and both samples were treated with alkyl DA-DA for 2 h. Click chemistry was used to detect the incorporated DA-DA in peptidoglycan. The samples were analyzed using Structured Illumination Microscopy. Images represent typical results from two independent experiments. **g.** The bacteria from the above experiment were subjected to FACS analysis. The mean fluorescence was used for the graph. n=3. ** Indicates p value <0.01. **h.** Renografin-purified EBs were incubated in axenic media containing only G6P or G6P together with Gln. After 24 h the bacteria were pelleted down and DNA was isolated to analyze the bacterial genome copy number. n=3. * Indicates p value <0.05.

Interestingly, significant ^13^C-labelling was also detected in Ala (17%) and the peptidoglycan precursor, diaminopimelate (DAP) (41%) (Fig. 1d). Whereas Ala was characterized by a high fraction of the M+3 isotopologue (*i.e.* representing a molecule with three ^13^C-labelled carbon atoms), the labelling pattern of DAP was more complex with the M+3, M+4 and M+7 isotopologues as the main species (Fig. 1d). The detected isotopologue composition of Ala can be explained by its formation from pyruvate derived from oxaloacetate *via* PEP catalyzed by PEP carboxykinase and pyruvate kinase and the decarboxylation of Asp although no gene is annotated for the decarboxylation of Asp in the genome of *Chlamydia* (scheme of the metabolic pathway in Supplementary Fig. 1d). The labelling pattern in DAP points at its formation from Asp and pyruvate contributing four and three labelled carbon atoms, respectively, and resulting in the detected M+3, M+4 and M+7 isotopologues species (Fig. 1d, supplementary Fig. 1d). In another experiment, we supplied Gln either labelled in the amino group or the amido group to the axenic culture. GC-MS analyses of Ala showed that the ^15^N-label was only transferred from the amino group of Gln (Supplementary Fig. 1b). Together, these results indicated that Gln serves as a major carbon source for the fueling of the partial TCA cycle of *Chlamydia* and as a source of amino groups for the generation of amino acids from alpha-keto acids via transamination, *e.g.* Glu, Asp and Ala, required for peptidoglycan biosynthesis.

Evidence for the synthesis of DAP in axenic medium was particularly intriguing since peptidoglycan synthesis has been reported to be restricted to the replicative phase and binary fission of *Chlamydia* and this has never been observed for *Chlamydia* outside their host cell ^2^. We therefore analyzed EBs from axenic cultures in the presence or absence of Gln by electron microscopy (EM). Surprisingly, incubation of EBs with Gln led to the morphological transition of a conversion intermediate between EBs and RBs, so-called intermediate bodies (IBs) ^9^ (Fig. 1e). Since only EBs are infectious, we tested the Gln-treated EBs/IBs for their capacity to infect HeLa cells. The infectivity of EBs incubated with Gln dropped to 20% compared to the control (Supplementary Fig. 1c), supporting the conclusion that Gln induces the developmental transition of EBs to a non-infectious form.

We then synthesized a clickable form of D-Ala to detect peptidoglycan in these bacteria, a method previously used to demonstrate chlamydial peptidoglycan synthesis in infected cells ^2^ (see Methods). Chlamydial particles incubated in the presence of Gln in this cell-free approach produced detectable peptidoglycan (Fig. 1f,g). Since peptidoglycan synthesis in *Chlamydia* has been shown to be associated with replication in host cells, we also investigated if *Chlamydia* shows replication in the presence of Gln by analyzing the genome copy number (Fig. 1h). Interestingly, chlamydial DNA amount almost doubled in the presence of Gln, indicating that Gln access initiates limited replication in *Chlamydia*.

### Chlamydial infection triggers altered glutamine metabolism in infected cells

Since our data suggested uptake of Gln as a central node for the interaction between host and bacterial metabolism, we investigated how *Chlamydia* infection influences the metabolism of primary human umbilical vein endothelial cells (HUVECs). LC-MS analysis of the supernatant from HUVECs infected with *Chlamydia* indicated increased depletion of most amino acids compared to uninfected control, indicative of their enhanced uptake by infected cells (Fig. 2a). Gln was one of the most depleted amino acids in the medium and Gln uptake was significantly increased in infected cells (Fig. 2a,b).

**Fig. 2:**
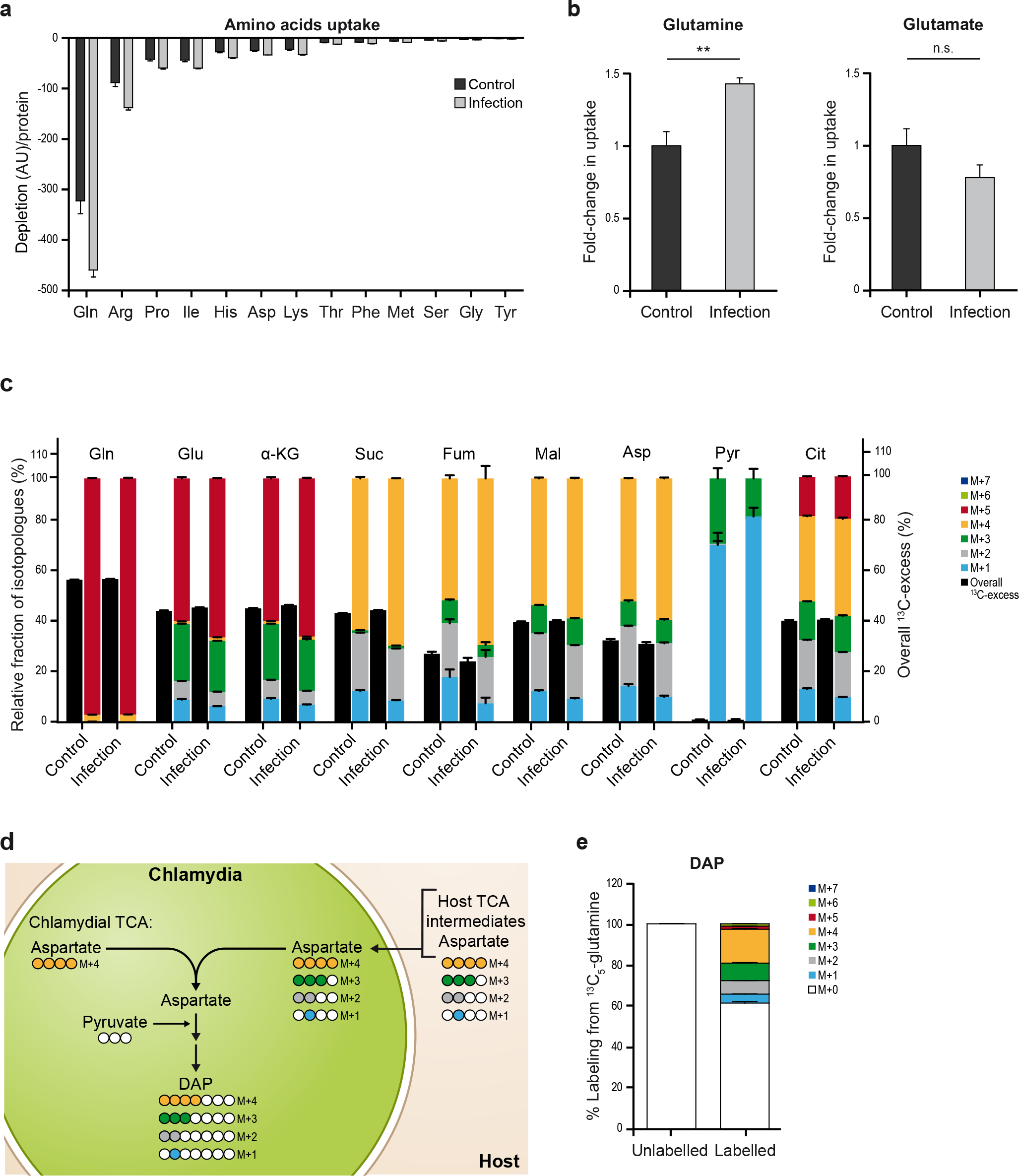
Chlamydial infection triggers altered glutamine metabolism in infected cells. HUVECs were either left uninfected or infected with *Chlamydia* at an MOI of 1 for 36 h. Cells or medium samples were extracted and metabolites were analyzed by LC-MS (see Methods). **a**. Depletion of amino acids from the culture medium from HUVECs with and without infection (only significant results are shown). **b**. Uptake of Gln and Glu from the medium by control and infected cells. n=3 * Indicates p value <0.05. **c**. Control or *Chlamydia* infected HUVECs were cultured in the presence of [U-^13^C_5_]Gln for 24 h. ^13^C-enrichments and isotopologue distributions of selected metabolites were determined by LC-MS. Data present the mean of three independent samples. **d**. Diagram outlining the synthesis of the D-aminopimelic acid (DAP) in *Chlamydia* from host or chlamydial derived metabolites. **e**. Mass isotopologue distribution of DAP extracted from infected host cells using acid hydrolysis. Data present the mean of three independent samples.

To further investigate the altered Gln metabolism in *Chlamydia*-infected cells, we performed an isotope labeling experiment by adding [U-^13^C_5_]-Gln to the culture medium that also contained unlabeled Gln at a ratio of 1:1 (i.e. 50% labelling). HUVECs were then infected with *Chlamydia* and incubated in this medium for 36 hours. LC-MS analysis revealed the labelling pattern of Gln, Glu, Asp and several TCA cycle metabolites (Fig. 2c). We also observed evidence for reductive carboxylation of Gln by the host cells as there was a substantial amount of the M+5 isotopologue of citrate. In contrast, Gln-derived labelling of pyruvate was almost absent (Fig. 2c), indicating that conversion of oxaloacetate to pyruvate by the enzymes of the gluconeogenic pathway is inactive in these cells under the culture conditions used. Interestingly, while the labelled fraction of Gln did not change upon infection, the M+5 labelled isotopologues of Glu and α-KG increased, indicating increased glutaminolysis in infected cells (Fig. 2c). Moreover, the M+4 isotopologues of succinate, fumarate, malate and aspartate also increased following *Chlamydia* infection, most likely due to enhanced entry of Gln-derived carbons into the TCA cycle (Fig. 2c).

To analyze if *Chlamydia* obtain building blocks for DAP biosynthesis from the host cell, we performed stable isotope labelling using [U-^13^C_5_]-Gln as described before and extracted bacterial cell wall components by acidic hydrolysis. LC-MS detection of DAP confirmed that glutamine-derived carbons are indeed incorporated into the bacterial metabolism to produce biomolecules essential for *Chlamydia* proliferation (Fig. 2d,e). Moreover, the high proportion of the M+4 isotopologue in DAP suggests that *Chlamydia* either takes up glutamine directly from the host cells and uses it to produce Asp via its truncated TCA cycle, or that glutamine is first converted by the host cell into metabolic intermediates that contain four ^13^C carbon atoms (*i.e.* succinate, fumarate, malate, oxaloacetate or Asp), which are then taken up by *Chlamydia* and used for the synthesis of DAP (Fig. 2c-e, diagram Fig. 2d and supplementary Fig. 2). Moreover, the high abundance of the M+1, M+2 and M+3 isotopologues in DAP also indicates that *Chlamydia* uses metabolic intermediates that are formed by the complete TCA cycle of the host cell. Production of these isotopologues depends on the second and third round of the TCA cycle, which requires the activity of citrate synthase, aconitase and isocitrate dehydrogenase present only in the host metabolism (see diagram in supplementary Fig.2). Together, these results clearly demonstrate that *Chlamydia-*infected cells increase uptake and metabolism of Gln to provide the pathogen with metabolic intermediates that are required for bacterial peptidoglycan production.

### *Chlamydia* infection stabilizes the proto-oncogene c-Myc

We next focused on how infected cells compensate for the increased Gln demand. RNA-Seq experiments were performed on infected and non-infected HUVEC cells and the gene expression profiles were compared. Gene Set Enrichment Analysis (GSEA) showed upregulation of MYC target genes in cells infected with *Chlamydia* (Fig. 3a and supplementary Fig.3a). *c-MYC*, the tightly regulated proto-oncogene is also known as a ‘master regulator’ of cellular metabolism ^10^, in particular mitochondrial glutamine metabolism ^11, 12^, and plays a role in apoptosis inhibition in *Chlamydia*-infected cells ^13^. We therefore examined the levels of c-Myc after infection with various species of *Chlamydia* in human and mouse cell lines. c-Myc levels were strongly increased in HUVEC cells already 12 hours post infection (hpi) with *C. trachomatis* and remained elevated up to 36 hpi (Fig. 3b). A similar effect was observed in different cell lines, including primary epithelial cells derived from human or mouse fimbriae (Fimb cells) (Supplementary Fig. 3b,c), and human osteosarcoma U2OS cells (Supplementary Fig. 3d), as well as upon infection of HeLa cells with different *Chlamydia* species (Supplementary Fig. 3e-g).

**Fig, 3:**
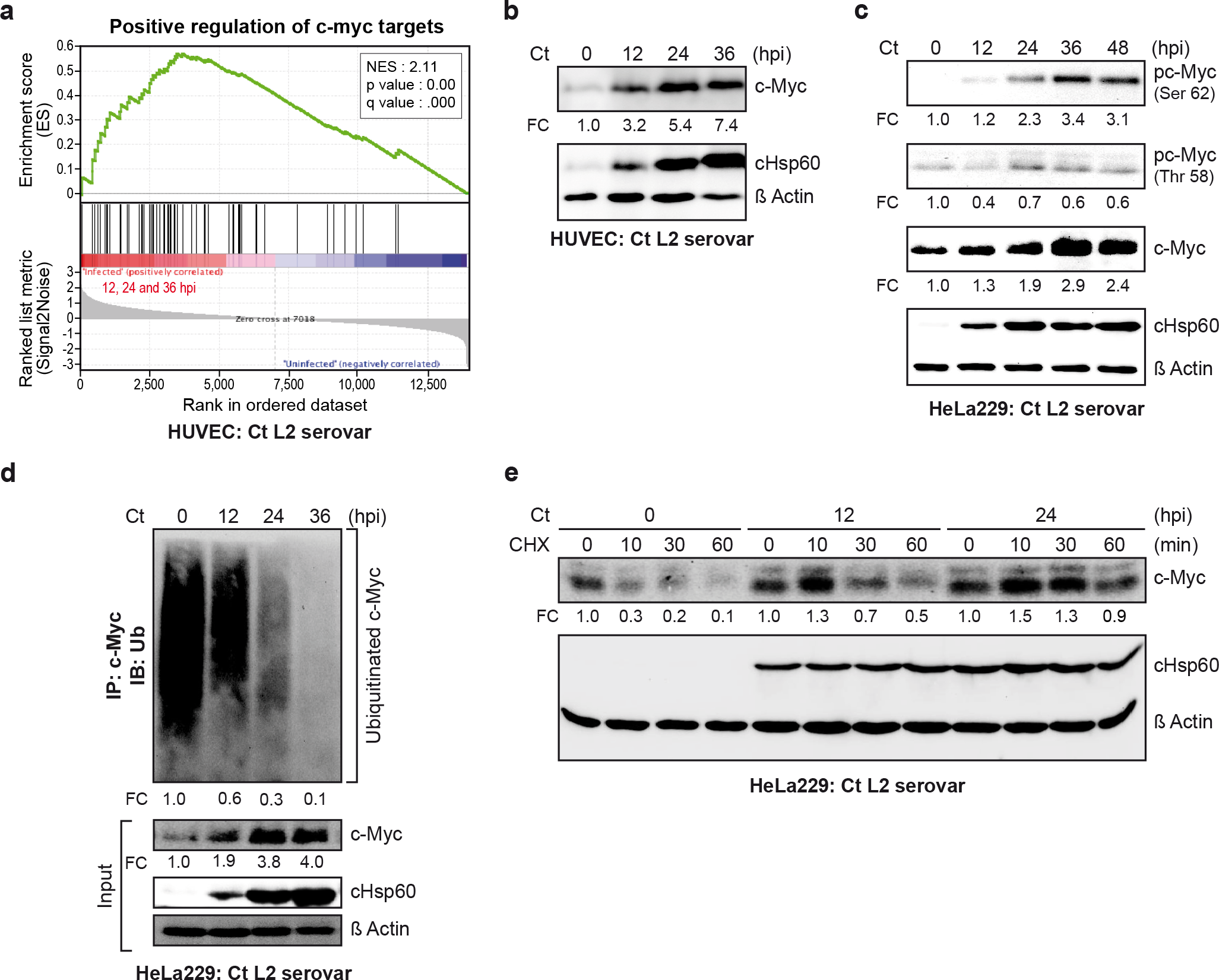
*Chlamydia* infection leads to upregulation and stabilization of the proto-oncogene c-Myc. **a.** Gene Set Enrichment Analysis (GSEA) ^40^ using the MSigDB Hallmark collection on genes sorted according to regulation between infected and uninfected conditions. Enrichment plot of MYC target gene set is shown as an example. **b.** HUVEC cells were infected with *Chlamydia* (Ct) at an MOI of 1 for different time intervals. The cells were lysed and c-Myc was detected by Western blot analysis. ß Actin served as the loading and chlamydial Hsp60 (cHsp60) as infection control. **c.** Hela229 cells were infected with *Chlamydia* at an MOI of 1 for different time points and the samples were analyzed using Western blotting for phosphorylated c-Myc (pc-Myc serine 62), pc-Myc (threonine 58), cHsp60 and ß Actin. **d.** c-Myc was immunoprecipitated from lysates of HeLa229 cells infected with *Chlamydia* for different time points. Ubiquitin was detected in the precipitate by immunoblotting. The input from the same experiment was probed against c-Myc, cHsp60 and ß Actin. **e.** Hela229 cells were either left uninfected or infected with *Chlamydia* at an MOI of 1 for 24 or 36 hpi. The cells were treated with the translation inhibitor cycloheximide (CHX) for 10, 30 or 60 min. The cells were then lysed and analyzed by Western blotting for c-Myc, cHsp60 and ß Actin.

We next investigated the potential mechanisms that could be involved in the observed induction of c-Myc protein in response to *Chlamydia* infection. Real-time PCR performed in *Chlamydia-*infected HeLa cells revealed a transient upregulation of c-Myc mRNA upon infection (Supplementary Fig. 3h). In addition to transcriptional regulation, phosphorylation of c-Myc at the conserved residues serine 62 (S62) and threonine 58 (T58) has been demonstrated to regulate c-Myc protein stability in response to mitogenic signaling ^14^. Phosphorylation of S62 transiently increases c-Myc stability whereas phosphorylation of T58 triggers dephosphorylation of S62 by protein phosphatase 2A (PP2A), ubiquitination by the SCF-Fbw7 E3 ligase, and proteasomal degradation ^15^. Interestingly, *Chlamydia* infection induced the phosphorylation of c-Myc at S62 but not at T58 (Fig. 3c). The level of c-Myc was increased in *Chlamydia-*infected cells and the protein accumulated in the nucleus (Supplementary Fig. 3i,j). In line with these findings, levels of ubiquitinated c-Myc were markedly decreased in *Chlamydia*-infected cells (Fig. 3d). We therefore determined whether *Chlamydia* infection altered the stability of the c-Myc protein. For this, cells were infected for different time points prior to the addition of the translation inhibitor cycloheximide. Whereas c-Myc levels rapidly declined in non-infected cells upon inhibition of translation, *Chlamydia*-infected cells retained c-Myc for extended periods (Fig. 3e), suggesting that c-Myc is stabilized upon infection.

### c-Myc is stabilized in *Chlamydia*-infected cells via MAPK and PI3K signaling pathways

*Chlamydia* infection is known to activate MAPK and PI3K pathways, which are critical for proliferation and growth of the pathogen ^16–18^. Activated Ras has been demonstrated to lead to the stabilization of c-Myc by inducing the phosphorylation of S62 through the MAPK/ERK pathway and by inhibition of T58 phosphorylation by phosphatidylinositol 3-kinase (PI3K) signaling (Fig. 4a) ^19^. Inhibition of MAPK or PI3K pathway using specific inhibitors indeed prevented the up-regulation of c-Myc protein levels and also attenuated the propagation of the bacteria in infected cells (Fig. 4b,c).

**Fig. 4:**
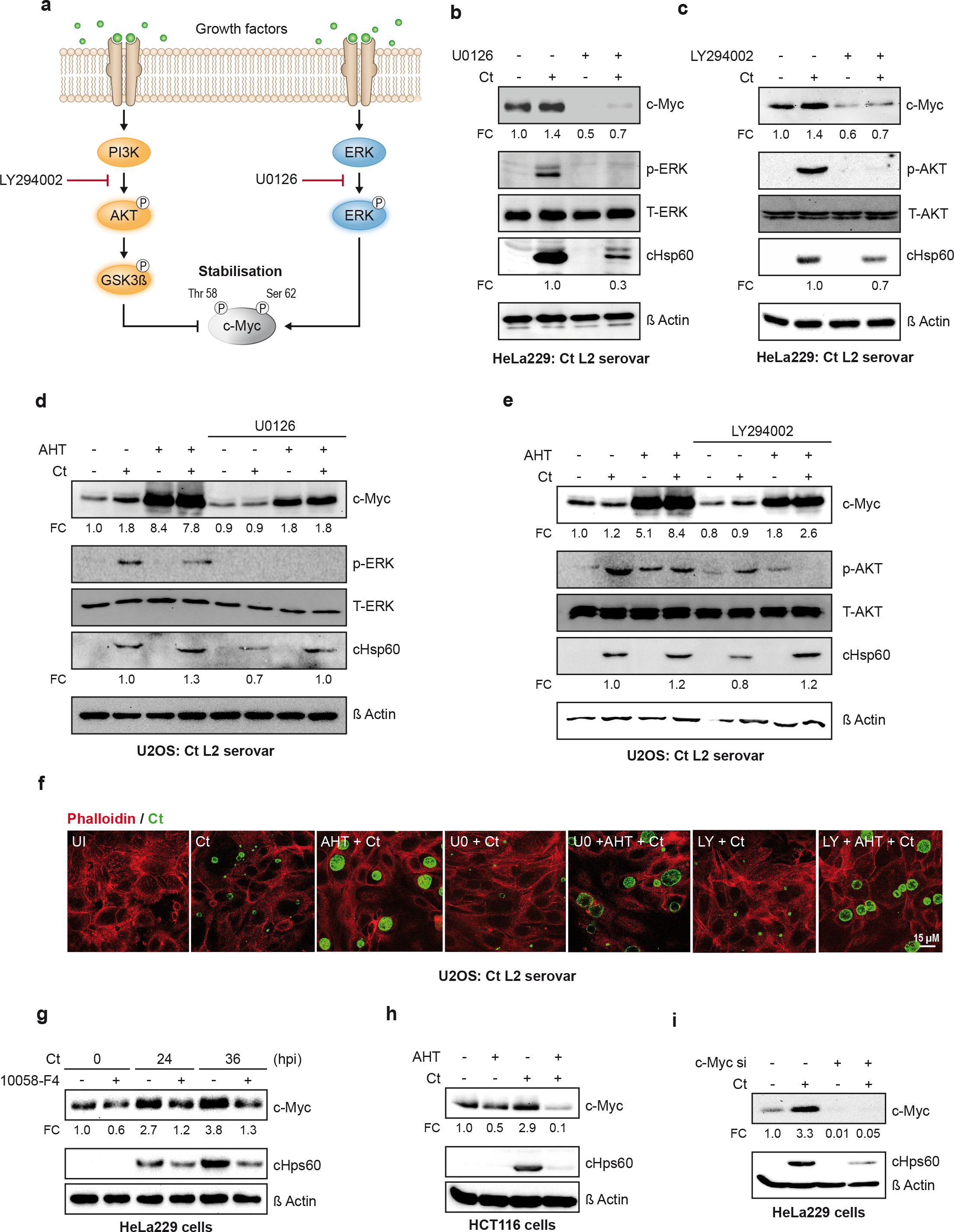
*Chlamydia* infection stabilizes c-Myc via the PI3K and MAPK pathway. **a.** Diagram showing the signaling pathway leading to c-Myc stabilization. **b./c.** HeLa229 cells were treated with the MAPK inhibitor, U0126 (30 μM) **(b)** or the PI3K inhibitor, LY294002 (30 μM) **(c)** for 4 h and infected with *Chlamydia* for 24 h. The cells were lysed and analyzed by Western blotting for c-Myc, pERK, T-ERK, pAKT, T-AKT as indicated. Detection of cHSP60 served as an infection and of ß Actin as a loading control. **d/e.** U2OS cells harboring an inducible c-Myc gene under the control of a Tet-inducible promoter were induced with AHT for 12 h and either treated with MAPK inhibitor U0126 **(d)**, or the PI3K inhibitor LY294002 **(e)** or left untreated as indicated. The cells were further infected with *Chlamydia* and analyzed by Western blotting for bacterial infection. **f.** The cells from the experiment shown in (d) and (e) were fixed and immunostained to detect chlamydial inclusions (green) and actin (phalloidin red). Images are representative for three independent experiments. **g.** HeLa229 cells were treated with the chemical c-Myc inhibitor 10058-F4. The cells were infected with *Chlamydia* for 24 or 36 hpi and then analyzed for *Chlamydia* infection by Western blotting (cHsp60). **h.** HCT116 cells engineered to express a shRNA to silence c-Myc expression under the control of an inducible Tet^on^ promoter were treated with AHT (1 μg/ml) for 24 h and then infected with *Chlamydia* for another 24 h. The cells were lysed and analyzed for *Chlamydia* infection by probing for cHsp60. **i.** siRNA pool was used to knock down c-Myc. After 48 h of transfection, the cells were infected with *Chlamydia* for 24 h. The cells were harvested and analyzed by Western blotting for chlamydial infection.

Intriguingly, *Chlamydia* growth was rescued in U2OS^Tet-On^ cells ^20^ upon anhydrous tetracyclin (AHT)-induced induction of c-Myc expression despite inhibition of the MAPK or PI3K pathway, respectively (Fig. 4d-f; AHT control Supplementary Fig. 4a), indicating that the anti-chlamydial activity of U0126 and Ly294002 is mediated by the down-regulation of c-Myc. Likewise, upon induction of c-Myc expression, the PI3K and MAPK inhibitors had no significant effect on the infectivity of *Chlamydia* (Supplementary Fig. 4b,c), demonstrating that *Chlamydia* develop normally under these conditions. The central role of c-Myc for chlamydial growth was further supported by experiments using the chemical inhibitor 10058-F4, a cell-permeable thiazolidinone that specifically inhibits c-Myc and transactivation of c-Myc target gene expression (Fig. 4g) and the depletion of c-Myc in HCT116 (Fig. 4h) or HeLa cells (Fig. 4i and supplementary Fig.4d).

### *Chlamydia* depends on host cell Gln uptake

Our gene set enrichment analysis (GSEA) on high throughput RNA sequencing data derived from control and *Chlamydia*-infected HUVECs cells revealed a strong influence of *Chlamydia* infection on the host metabolite transporter and glutamine metabolic pathway (Supplementary Fig. 5a,b). To understand if Gln is necessary for the intracellular growth of *Chlamydia*, we used Gln-depleted media for infection studies. *Chlamydia* failed to replicate in HeLa229 and primary human Fimb cells cultured in medium without Gln (Fig. 5a,b and supplementary Fig. 5c,d). This was accompanied by a failure to induce c-Myc expression in infected cells (Fig. 5a). Bacteria harvested from these cultures and transferred to fresh cells in the presence of Gln failed to initiate an infection, demonstrating that no infectious progeny were produced in the absence of Gln (Fig. 5b, supplementary Fig. 5e).

**Fig. 5:**
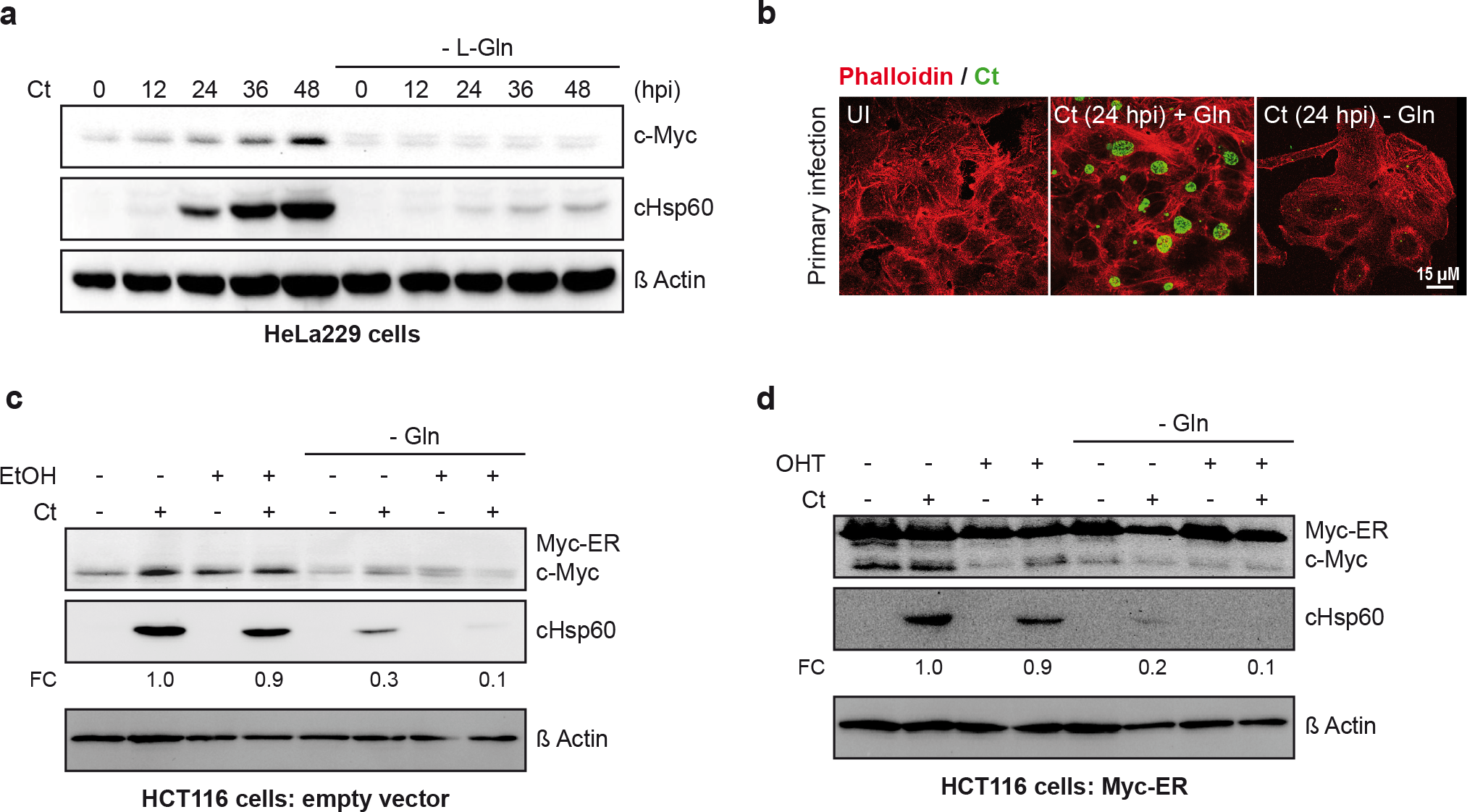
Glutamine is a limiting metabolite for chlamydial intracellular growth. **a.** HeLa229 cells were grown in basic formulation of DMEM containing 1g/L D glucose. The cells were treated with or without glutamine (2 mM) and infected with *Chlamydia* for different time points. The cells were lysed and analyzed for chlamydial growth using Western blotting to detect chlamydial Hsp60, c-Myc and ß Actin. **b.** The cells from (**a)** were fixed and immunostained for chlamydial inclusions (green) and actin (phalloidin, red). **c/d.** HCT116 were grown in basic formulation of DMEM containing 1g/L D glucose, with or without L-glutamine. HCT116 expressing 4-hydroxytamoxifen (OHT)-inducible Myc-ER or the vector control were treated overnight with OHT (4100 nM) **(d)** to activate Myc-ER or ethanol (1 μl) as control **(c).** The cells were then infected with *Chlamydia* at an MOI of 1 for 24 h and analyzed for bacterial load by Western blotting.

It has been shown recently that c-Myc can function in a glutamine sensing pathway since depletion of glutamine causes the rapid depletion of c-Myc via a mechanism dependent on the 3’ UTR of the gene ^21^. To rule out that Gln depletion affects *Chlamydia* growth indirectly via c-Myc downregulation, we used HCT116 cells expressing the MYC-ER fusion protein from a construct lacking the 3’UTR, which allows the restoration of c-Myc expression in Gln-deprived cells. However, Gln depletion severely attenuated the growth and development of *Chlamydia* in this setting (Fig. 5c,d), indicating that c-Myc cannot rescue chlamydial growth in the absence of Gln.

### *Chlamydia* infection reprograms host cell metabolism by inducing glutamine uptake and catabolism

The unexpected result of an exclusive uptake of Gln by *Chlamydia* observed in axenic cultures suggested that Gln is also a central host amino acid used by *Chlamydia* for peptidoglycan biosynthesis and for complementing the partial TCA cycle. The question therefore was how the infected cells deal with these increased levels of Gln required by fast replicating *Chlamydia.* We therefore investigated the regulation of genes related to glutamine uptake and catabolism during infection. Interestingly, GSEA revealed a set of glutamine transporter genes including SLC1A5/ASCT2 ^22^ being upregulated during infection (Fig. 6a supplementary Fig. 6a). Furthermore, the mRNA for glutaminase (GLS1), one of the enzymes converting Gln to Glu (Fig. 6b), was also strongly induced upon *Chlamydia* infection (data not shown).Both, SLC1A5/ASCT2 and GLS1 have been shown to be c-Myc target genes induced to enhance glutamine uptake and catabolism in cancer cells ^11, 23^.

**Fig. 6:**
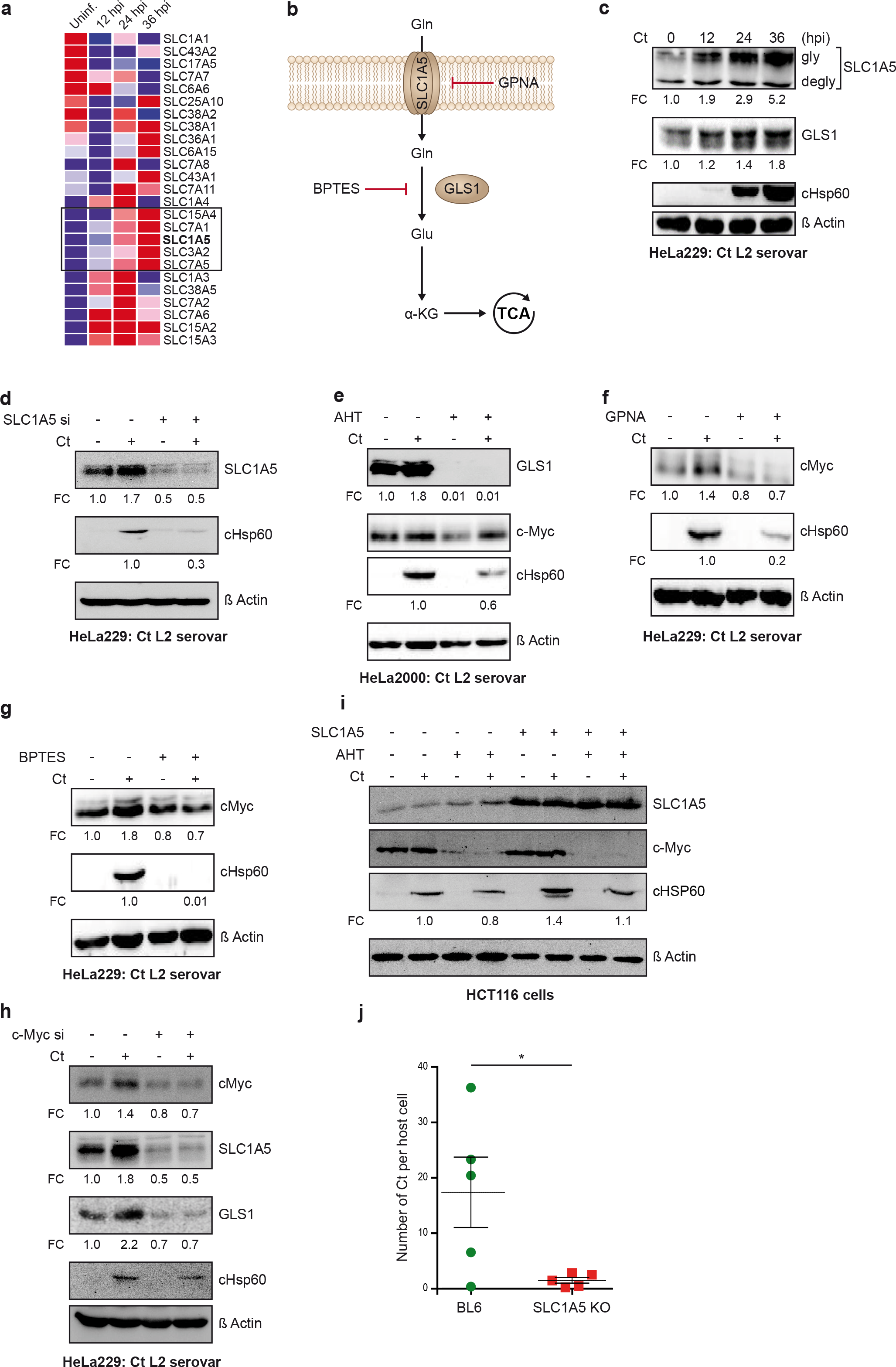
*Chlamydia* metabolically reprograms the host cell by up regulating SLC1A5 and glutaminase. **a.** HUVEC cells were infected with *Chlamydia* for different time points and the total RNA was isolated and analyzed by RNA Seq. The heat map from an example gene set enriched in infected cells (Reactome Amino Acid and Oligopeptide SLC Transporters) generated with GSE analysis is shown. Significantly upregulated transporters are boxed. **b.** Diagram showing Gln uptake and shuttling into the TCA cycle and the targets of the inhibitors used to block Gln uptake and metabolism. **c.** HeLa229 cells were infected with *Chlamydia* at an MOI of 1 for different time points. The cells were lysed and SLC1A5 and GLS1, cHsp60, and ß Actin were detected by Western blot analysis. **d.** siRNA pool was used to knock down SLC1A5. 48 h after the transfection, the cells were infected with *Chlamydia* at an MOI of 1 for 24 h. The cells were lysed and further analyzed by Western blot. **e.** HeLa229 cells expressing a shRNA to silence GSL1 expression under the control of a Tet^on^ promoter were induced with AHT for 7 days to completely deplete GLS1. These cells were then infected with *Chlamydia* for 24 h and the cell lysate were used to determine the levels of GLS1, cHsp60, and ß Actin. **f.** HeLa229 cells were treated with the SLC1A5 specific inhibitor GPNA for 4 h and infected with *Chlamydia* at an MOI of 1 for different time points. The cells were lysed and analyzed by Western blotting. **g.** HeLa229 cells were treated with the glutaminase inhibitor BPTES for 4 h and infected with *Chlamydia* at an MOI of 1 for different time points. The cells were lysed and Western blot analysis was performed to detect c-Myc, cHsp60, ß Actin. **h.** HeLa229 cells were transfected with siRNA against c-Myc. 48 hpi, the cells were analyzed by Western blotting as indicated. **i.** HCT116 Tet^on^ sh-c-Myc cells were either left untreated or treated with AHT (1μg/ml) for 24 h followed by overexpression of SLC1A5. The cells were then infected with *Chlamydia* at MOI of 1 for another 24 h and analyzed by Western blotting. **j.** C57BL6 or SLC1A5 knock out mice were infected with *Chlamydia*. The mice were sacrificed seven days post infection and the copy number of *Chlamydia* was calculated using RT-PCR. * Indicates p value <0.05.

Western blots of control and *Chlamydia*-infected samples revealed a 5.2-fold up-regulation of SLC1A5 and a 1.8-fold up-regulation of GLS1 (Fig. 6c). Furthermore, depletion of SLC1A5 or GLS1 using short interfering RNA (siRNA) or shRNA (Fig. 6d,e) or blocking the activity of SLC1A5 or GLS1 using the specific chemical inhibitors GPNA and BPTES (Fig. 6f,g) drastically reduced replication of the pathogen. Since SLC1A5 and GLS1 are transcriptional targets of c-Myc ^11, 24^, we knocked down c-Myc, which resulted in strong depletion of both proteins and also blocked chlamydial growth (Fig. 6h).

Most interestingly, when we depleted c-Myc in cells overexpressing SLC1A5, we could retain *Chlamydia* growth during the primary infection but the progeny failed (Fig. 6i and supplementary Fig.6b). This result supports a role of Gln in establishing a primary infection in line with our observation that Gln triggers the EB-RB transition (Fig. 1e). However, c-Myc regulated host cell functions, for example the production of TCA cycle intermediates, may be essential for the full development of *Chlamydia*. Taken together, these data demonstrate a critical role of c-Myc and its transcriptional targets SLC1A5 and GLS1 in the reprogramming of the Gln metabolism of host cells to promote *Chlamydia* replication and development.

Since our results showed that Gln is absolutely critical for *Chlamydia*, we asked if reprogramming of Gln metabolism would also affect chlamydial development *in vivo*. Recently, SLC1A5 knockout mice have been generated (Supplementary Fig. 6c), which show no major phenotype ^25^. Interestingly, trans-cervical infection of these mice revealed a significant growth defect of *Chlamydia* in the absence of SLC1A5 compared to control (Fig. 6j), suggesting that limiting the uptake of this non-essential amino acid could serve as a novel therapeutic approach against *Chlamydia* infection.

## Discussion

After the completion of the developmental cycle EBs released from the host cell have to avoid the transition to RBs since only EBs are capable of entering fresh host cells and initiate a new generation of progeny. The longstanding view that EBs are metabolically inactive has recently been revised since EBs kept in axenic medium with G6P, a metabolite available to *Chlamydia* only inside cells, generate ATP via glycolysis ^7^. However, despite this active metabolism, EBs do not initiate the conversion to RBs under these conditions. Here we provide evidence that Gln is the key metabolite that initiates peptidoglycan synthesis and the EB to RB transition. The concentration of Gln in uterine fluid is as low as 0.13 mM ^26^, however, inside cells Gln reaches levels of 2 to 30 mM ^27^. The findings that *Chlamydia* selectively take up Gln for peptidoglycan synthesis and the pattern of Gln concentrations outside and inside of cells support our hypothesis of Gln as the metabolic trigger for EB to RB transition.

Our finding that L-glutamine serves as a crucial amino acid for the replication of *Chlamydia* corroborates previous findings ^28, 29^. Many of the glutamine-derived intermediates in *Chlamydia* serve as precursors for peptidoglycan biosynthesis (Fig. 1d and supplementary Fig. 1d) underlining the central role of glutamine for the chlamydial metabolism. Chlamydiae do not form a peptidoglycan sacculus typical for other Gram negative bacteria but only assemble peptidoglycan rings in the mid-cell of actively dividing RBs.. We detected the accumulation of peptidoglycan for the first time also in *Chlamydia* outside of host cell but only in the presence of Gln (Fig. 1f.g), where we also observed a change in the morphology of the EBs (Fig. 1e). Peptidoglycan accumulated in or close to the bacteria and was not assembled as a ring which could indicate that the crosslinking of the peptidoglycan disaccharide pentapeptide did not occur in this setup. The lack of lipids, which they normally acquire from the host for stabilizing the peptidoglycan ring, might explain this observation. The increase in the copy number of the bacterial genomes (Fig. 1h), however, indicated a start of the replicating machinery. *Chlamydia* is an auxotroph for nucleosides and may use the stored metabolites to initiate replication in axenic culture.

*Chlamydia* like many other obligate intracellular bacteria replicate in differentiated cells that run the reduced metabolism of non-dividing cells. A single chlamydial particle can produce up to ∼ 1500 progeny in one infection cycle in a single cell. Since all metabolites of replicating *Chlamydia* originate from the host cell, the metabolism of the cell has to drastically change to meet the requirements of the infected cell. Glutamine is one of the most abundant amino acids in serum and fast-growing cells take up glutamine to support anabolic metabolism at multiple nodes, for example by using glutamine for protein synthesis, as a precursor for the synthesis of other amino acids, as a nitrogen donor for nucleotide biosynthesis, for the production of glutathione and as a substrate for TCA cycle anaplerosis. We show here that *Chlamydia* depend on the direct uptake of glutamine as well as on glutamine-derived host metabolites and therefore reprograms the host cell by stabilizing the central metabolism regulator c-Myc. The profile of c-Myc regulated genes and the dependence of the replicating *Chlamydia* on glutaminolysis parallels the reprogramming of dormant cells to fast proliferating and in particular tumor cells ^30^. c-Myc promotes glutamine uptake and glutaminolysis by increasing the expression of the glutamine transporters ASCT2/SLC1A5 and SNAT5/SN2 and of glutaminase (GLS) ^12^. We demonstrate here for the first time that bacteria induce host cell glutamine uptake and glutaminolysis for their replication. In addition, chlamydial growth was severely affected in genital infections of ASCT2/SLC1A5 knockout mice. This first demonstration of a dominant role of host cell glutamine reprogramming in an *in vivo* infection at all was surprising since this knockout mouse has no other major phenotype ^25^. Gln is channeled into different anabolic and catabolic pathways by glutaminolysis, generating Glu and subsequently α-KG, both of which are essential for infection and replication of several viruses ^31–36^. However, the relevance of glutamine reprogramming for viral infection still has to be shown *in vivo*.

In addition to glutamine transporters and glutaminase, we find other prominent c-Myc-regulated amino acid transporters like SLC43A1, SLC7A11, SLC15A4, SLC7A1, SLC3A2 and SLC7A5 are also upregulated during infection (Fig. 6a and supplementary Fig. 6a), pointing to a c-Myc-dependent central reprogramming of the amino acid supply in the infected cell. This global reprogramming of infected host cells by c-Myc stabilization may explain our intriguing finding of only a partial rescue of chlamydial infection by overexpression of SLC1A5 in c-Myc-depleted cells. The provision of increased glutamine levels is essential, but not sufficient to permit the complex cycle of chlamydial replication and development.

We show here the central role of glutamine in peptidoglycan biosynthesis, the initiation of the EB to RB transition, the complementation of the bacterial and host cell TCA cycle, and as nitrogen source. *Chlamydia* reprograms the glutamine metabolism of the host cell through c-Myc-controlled metabolic pathways to support its intracellular lifestyle. The central addiction of *Chlamydia* to host glutamine is reminiscent of cellular proliferation, malignant transformation and a hallmark of cancer cells which use extracellular glutamine to fulfill the metabolic demands of producing cell mass. Glutamine addicted tumor cells and, as we show here, *Chlamydia*-infected cells are highly sensitive to pharmacological interference with glutamine metabolism. Current approaches to target c-Myc or glutamine metabolism as target for innovative cancer therapy may prove to be also efficient in treating *Chlamydia* infection.

## Methods

### *Chlamydia* serovar used in the study

*Chlamydia trachomatis* (serovar L_2_/434/Bu and D) were used in this study. Some experiments were also performed using *Chlamydia muridarium* and *Chlamydia pneumoniae*. *Chlamydia* was prepared as previously published. Briefly, *Chlamydia* were grown in HeLa229 cells (ATCC^®^ CCL2.1^™^) at an MOI (multiplicity of infection) of 1 for 48 hour (h). The cells were lysed using glass beads (15 mm) for 3 minutes (min) and centrifuged at 2000 g for 10 min to remove the cell debris. The supernatant containing bacteria was collected and centrifuged at 24,000 g for 30 min at 4°C. The pellet was washed and resuspended in SPG buffer (0.25 M sucrose/10 mM sodium phosphate/5 nM glutamic acid), aliquoted and stored in −80°C. *Chlamydia* EBs and cell lines used in the study were verified to be free of Mycoplasma contamination via PCR. The bacteria were titrated to have an MOI of 1 and were used further in the experiments. After 1 hpi the media was replaced with fresh RPMI containing 5% FCS, infected cells were cultured in 37°C and 5% CO_2_.

### Culture of *C. trachomatis* in axenic medium

*Chlamydia trachomatis* L2 was propagated in HeLa229 cells, isolated, purified and incubated in different axenic media as previously described (Mehlitz et al., 2016). In brief, HeLa229 cells were seeded in T175 flasks and infected at an MOI of one. Forty-eight hours post infection, cells were scraped off, disrupted with glass beads and EBs were purified using 60-20% Renografin gradient (Meglumin diatrizoate (Sigma-Aldrich M5266), Sodium diatrizoate hydrate (Sigma-Aldrich S4506), Sodium citrate hydrate (Applichem A4522), EDTA (Servca 11280) add to 50 ml HBSS (Gibco 14025-050) pH-7.4, sterile filtered in 0.2 μm filter and stored at 4°C). EBs were resuspended in axenic media (basic DMEM (Sigma-Aldrich) supplemented with sodium bicarbonate (44 mM), phenol red (42 µM) and glucose-6-phosphate (0.5 mM) and incubated for respective time points at 37°C. According to the experimental set-up, 1mM of L glutamine / L-glutamate / pyruvate / α-ketoglutarate / [U-^13^C_5_]glutamine / [^15^N-amine]glutamine or [^15^N-amide] glutamine was added to the media. After incubation, samples were centrifuged for 30 min at 21,500 g at 4°C and supernatant was transferred to a new tube. Supernatant and pellets were heat-inactivated (10 min at 90°C) and stored at −80°C for further analysis.

### Substrate uptake analysis from axenic culture

#### a. α-ketoglutarate

0.2 mL of the supernatant from the axenic medium was spiked with 20 µl of a 5 mM norvaline solution (internal standard) and dried under N_2_ flux. The residue was treated with 50 µl methoxyamine in pyridine (20mg/ml) at 40°C for 90 min, and subsequently with 50 µl (N-(tert-butyldimethylsilyl)-N-methyl-trifluoroacetamide containing 1% tert-butyldimethylsilylchloride) (MTBSTFA) at 70°C for 30 min. This solution was taken for analysis.

#### b. Glutamate and glutamine

0.1 mL of the supernatant from the axenic medium was spiked with 20 µl of a 5 mM non-labeled glutamate or glutamine solution (internal standard) and dried under N_2_ flux. The residue was treated with 50 µl acetonitrile and 50 µl MTBSTFA at 70°C for 30 min and taken for analysis.

### Isotopologue profiling with *Chlamydia* from axenic culture

Bacterial pellets were suspended in 1mL of methanol and were mechanically disrupted using a ribolyser (3×20 sec 6.5 m/s). Afterwards the solution was centrifuged (10,000 g for 20 min, 4°C). This procedure was performed twice. The supernatants were combined and then dried under N_2_ flux. The residue was treated with 50μL of MTBSTFA and 50 μL of water free acetonitrile at 70°C for 30 min. The *tert*-butyl dimethylsilyl (TBDMS)-derivatives of amino acids and other metabolites were then analyzed by GC/MS. The residual cell debris after centrifugation was subjected to acidic hydrolysis as described earlier ^37^ and protein bound amino acids as well as diaminopimelate (DAP; retention time, 24.48 min; m/z 589) were analyzed as TBDMS derivatives.

### GC/MS conditions

All derivatives mentioned above were analyzed by GC-MS using a GCMS-QP 2010 Ultra spectrometer (Shimadzu, Duisburg, Germany) equipped with a EquityTM-5, fused silica capillary column, 30 m × 0.25 mm × 0.25 m film thickness. All data were collected using μm film thickness. All data were collected using LabSolution software *(Shimadzu)*. The samples were analyzed three times as technical replicates. The overall ^13^C excess (mol-%) and the relative contributions of isotopomers (%) were computed by an Excel-based in-house software package according to ^38^.

### TBDMS-derivatives of polar metabolite mixtures

The column was first developed at 100°C for 2 min, then using a gradient of 3°C min^−1^ to 234°C, followed by 1°C min^−1^ to 237°C and 3°C min^−1^ to 260°C. Finally, the column was heated at a gradient of 10°C min^−1^ to a final temperature of 320°C where it was hold for 2 min.

### Analysis of TBDMS-amino acids and DAP

The column was first developed at 150°C for 3 min, then using a gradient of 7°C min^−1^ to 280°C where it was hold for 5 min.

### Transmission electron microscopy

Chlamydial EBs were incubated with the axenic medium with G6P and with or without Gln. The bacteria pellet was fixed with 2.5% glutaraldehyde (50 mM sodium cacodylate (pH□7.2), 50□mM KCl, 2.5□mM MgCl_2_) at room temperature. The cells were incubated for 2□h at 4□°C with 2% OsO4 buffered with 50□mM sodium cacodylate (pH□7.2), washed with distilled H_2_O and incubated overnight at 4□°C with 0.5% uranyl acetate (in distilled H_2_O). The cells were dehydrated, embedded in Epon812 and ultrathin-sectioned at 50□nm. Sections were stained with 2% uranyl acetate in ethanol followed by staining with lead citrate and analysed in a Zeiss EM10 microscope (Zeiss). Electron micrographs were processed using ImageJ (Fiji).

### Click chemistry

Click chemistry was performed as described in ^2^. Axenic culture with purified EBs from *Chlamydia* was fed with ADA-DA (10μM). Clickable’ Alexa Fluor 532-azide and Click-iT® Cell Reaction Buffer Kit were purchased from Invitrogen.

### Copy number of *Chlamydia* genomes in axenic culture

*Chlamydia* were grown in HeLa229 for 48 h. The cells were lysed and the EBs were purified by renografin gradient separation as explained above. The bacteria were pooled by centrifugation and re-suspended in axenic medium without Gln. The re-suspended *Chlamydia* were split into two aliquots and Gln was added into one of them and incubated at 35°C for 24 h. The bacteria were further pelleted and DNA was isolated using DNAzol reagent (Thermo Fisher Scientific). Quantitative PCR was used to enumerate *Chlamydia* genome copy number. The following primers were used for amplifying the *C. trachomatis lytA* gene that was cloned into the vector: fwd, 5′-TCTAAAGCGTCTGGTGAAAGCT-3′ and rev, 5′-GAAATAGCGTAGTAATAATACCCG-3′. Data were analyzed by using the Step One Plus software package (Applied Biosystems). GraphPad Prism 7 was used to generate the graph.

### Cell culture and transfection

HeLa229 were used for propagating bacteria and for basic experiments. Epithelial cells isolated from human fimbriae (Fimb cells), U_2_OS (ATCC^®^ HTB-96^™^), HCT116 (ATCC^®^ CRL-247^™^) and HUVECs (ATCC^®^ CRL-1730^™^) were also used in the study. HUVECs were used in the high throughput RNA sequencing. All cell lines were tested negative for mycoplasma contamination via PCR. HeLa229 and human Fimb cells were grown in RPMI1640 + GlutaMAX^TM^ (Gibco^TM^ 72400-054) with 10% heat inactivated FCS (Sigma-Aldrich F7524). U_2_OS and HCT116 cells were cultured in DMEM (Sigma-Aldrich D6429) with 10% heat inactivated FCS. HUVECs were cultured in Medium 200 (Gibco^TM^ M200500) containing 1x LSGS (Gibco^TM^ S00310). For glutamine deprivation experiments, the cells were first seeded in RPMI1640 + GlutaMAX^TM^ or DMEM, high glucose (Sigma-Aldrich D6429). The following day the medium was changed to the basic formulation of DMEM (Sigma-Aldrich D5030) supplemented with 5% dialyzed FCS (Sigma-Aldrich F0392), 1 or 4.5 g/L D-glucose (for HeLa229/ U_2_OS or human Fimb respectively) and varying concentrations of L-glutamine (according to experimental setup).

Cells were transfected with plasmid DNA at a confluency of 60% with Polyethylenimine (PEI) or X-tremeGENE^TM^ HP DNA transfection reagent (Roche) and OptiMEM transfection medium (Gibco) in 5% FCS medium. After 5 h, transfection medium was replaced by fresh RPMI supplemented with 5% FCS medium. The plasmids used in the study are described in the supplementary file. siRNA against SLC1A5 (sc-60210) and c-Myc (sc-29226) was obtained from Santa cruz Biotech.

### Metabolic profiling

For this study HUVECs were seeded in triplicates, either uninfected or infected with *C. trachomatis* serovar L2 for 36 hours. After the respective time medium was collected, snap frozen in liquid nitrogen, and the cells were washed with ice cold 154 mM ammonium acetate (Sigma) and snap frozen in liquid nitrogen. The cells were harvested after adding 480 μl cold MeOH/H2O (80/20, v/v) (Merck) to each sample containing Lamivudine (Sigma) standard (10 µM). The cell suspension was collected by centrifugation and transferred to an activated (by elution of 1 mL CH3CN (Merck)) and equilibrated (by elution of 1 mL MeOH/H2O (80/20, v/v)) RP18 SPE-column (Phenomenex). The eluate was collected and evaporated in SpeedVac and was dissolved in 50 μL CH3CN/5mM NH4OAc (25/75). Each sample was diluted 1:1 (cells) or 1:5 (medium) in CH3CN. 5 µl of sample was applied to HILIC column (Acclaim Mixed-Mode HILIC-1, 3 µm, 2.1 * 150 mm). Metabolites were separated at 30°C by LC using a DIONEX Ultimate 3000 UPLC system (Solvent A: 5mM NH4OAc in CH3CN/H2O (5/95), Solvent B: 5 mM NH4OAc in CH3CN/H2O (95/5); Gradient: linear from 100% B to 50% B in 6 min, followed by 15 min const. 40% B). MS-Analysis was done on a Thermo Scientific QExactive instrument in positive and negative mode. Peak determination and semi-quantitation was performed using TraceFinder™ Software. For determination of protein content for the data normalization, BCA assay (Thermo Fisher Scientific) was performed. The pellet of the cell samples was dried, resuspended in 0.2 M sodium hydroxide (Roth), boiled for 20 min at 95°C and absorbance was measured at 550 nm. Prism GraphPad was used for statistical analysis.

### Western blotting and antibodies

Lysates for Western blot analysis were prepared by directly lysing cells in SDS sample buffer (62.5 mM Tris, pH 6.8, 2% SDS, 20% glycerol and 5% ß-mercaptoethanol) in ice. Western blot analysis was performed as described in ^39^. Briefly protein samples were separated in the 6-12% SDS-PAGE (Peqlab) and transferred to a PVDF membrane (Roche) in a semidry electroblotter (Thermo Fisher Scientific). The membrane was further blocked in tris buffer saline containing 0.05% Tween20 and 5% bovine serum albumin or dry milk powder. The primary antibody against c-Myc (Y69: ab-32072), pc-Myc Thr58 (ab-28842) and pc-Myc Ser62 (ab-51156) SLC1A5 (ab-84903) glutaminase (ab-156876) was purchased from Abcam. The T-ERK (cs-9180), pERK (cs-9106), T-AKT (cs-9272), pAKT Ser473 (cs-9271), were obtained from Cell Signaling. Chlamydial HSP60 (sc-57840),) and anti-ubiquitin (sc-8017) antibody was purchased from Santa Cruz Bioscience and ß Actin antibody from Sigma (A5441). Proteins were detected with secondary antibodies coupled with HPR (Santa Cruz Bioscience) using ECL system (Pierce) and Intas Chem HR 16-3200 reader. Quantification of blots was done by FIJI (ImageJ) software.

### Immunoprecipitation

Uninfected and *Chlamydia*-infected (MOI 1) HeLa229 cells were lysed using denaturing buffer (RIPA lysis buffer: 50mM Tris-HCl pH-7.5, 150mM NaCl, 1% Triton-X100, 1% NP-40, 0.1% SDS, 10% glycerol containing Complete protease inhibitor cocktail (Roche) and MG-132, proteasome inhibitor) to prevent co-precipitation of interacting partners of c-Myc. Lysates from 7 × 10^6^ cells were prepared as described before and incubated with 3 µg anti-c-Myc antibody for 1 h at 4°C followed by incubation with protein G magnetic beads (Dynabeads, Thermo Fisher Scientific) for 2 h at 4°C. The samples were washed several times and eluted by addition of 2x SDS-sample buffer and heating to 94°C. Samples were separated with SDS-PAGE and visualized by immunoblotting after probing against anti ubiquitin antibody.

### Nuclear-cytoplasmic isolation

HeLa229 cells were plated in 150 mm dishes and either left uninfected or infected with *Chlamydia* (MOI 1) for the mentioned period of time. The cells were washed with ice cold PBS. The cells were scraped into a falcon. The cells were centrifuged and re suspended in buffer containing 10 mM Hepes-KOH pH 7.9, 10 mM KCl, 1.5 mM MgCl2, 0.5 mM DTT, 0.05% NP-40, protease inhibitors and incubated for 20 min in ice. The cells were then homogenized with a dounce homogenizer (10 strokes). The cells were centrifuged at 4000 RPM for 5 min at 4°C, the supernatant containing cytoplasmic proteins was collected and lysed with 2x SDS-sample buffer. The pellet was resuspended in buffer containing 20 mM Hepes-KOH pH 7.9, 400 mM NaCl, 1.5 mM MgCl2, 0.2 mM EDTA, 15 % glycerol, 0.5 mM DTT, protease inhibitors and incubated for another 20 min and vortexed. Further the sample was centrifuged at 14000 RPM for 10 min at 4°C, the pellet containing the nuclear extract was lysed in 2x SDS-sample buffer and analyzed using Western blot.

### Immunofluorescence analysis

The immunostaining was performed as described earlier. Briefly, HeLa229 and HUVEC cells were grown on cover slips and infected with the indicated *C. trachomatis* strain at an MOI 1 for indicated time points. The cells were washed with PBS, fixed with 4% PFA/Sucrose and permeabilized with 0.2% Triton-X-100/PBS for 30 min. Samples were blocked with 2% FCS/PBS for 1 h. All primary antibodies were incubated for 1 h at room temperature. Primary antibodies were used in the following dilutions in 2% FCS/PBS: anti-HSP60 (1:500), c Myc (1:200). Samples were washed three times and incubated with a Cy2-/Cy3-/Cy5-conjugated secondary antibody for 1 h in the dark. The cells were mounted on microscopic slide using Mowiol. Slides were air dried for at least 24 h and examined using Leica DM2500 fluorescence microscope, the images were analyzed using LAS AF and Image J software.

### Inhibitor studies

Hela229 cells were grown in the RPMI1640 +GlutaMAX^TM^ or in the basic DMEM medium supplemented with 5% dialysed FCS and 1 g/L of D-Glucose and 4mM L-glutamine. The cells were treated with MAPK inhibitors LY297004/UO126 (30μM) or c-Myc inhibitor (10058-F4: Sigma-Aldrich)/ the glutaminase inhibitor (BPTES (5μM): Sigma-Aldrich) or the SLC1A5 inhibitor (GPNA (5mM): Sigma-Aldrich) for 2 h and the cells were infected with *Chlamydia* at an MOI 1 for the respective time point. The vehicle used for each inhibitor was used in the appropriate concentration. The cells were lysed and analyzed using Western blotting.

### RNA Sequencing and NGS Data Analysis

Libraries for RNA Seq were generated using NEBNext® Ultra™ RNA Library Prep Kit for Illumina® with 12 PCR cycles for amplification and sequenced on Illumina NextSeq500 platform. FASTQ generation was carried out using CASAVA and the quality check was performed using FastQC. Reads from FASTQ files were aligned to hg19 genome using Bowtie2 and differential gene regulation calculation was carried out based on edgeR algorithm. Counts of reads normalized to library size were used for carrying out Gene Set Enrichment Analysis (GSEA) using weighted parameters on C2, C5 and Hallmark gene sets downloaded from MSigDB database ^40^.

### Real Time PCR

RNA was isolated from uninfected and *Chlamydia*-infected HeLa229 cells using RNA easy kit (Qiagen, Germany). RNA was reverse transcribed using a Revert Aid First Strand synthesis Kit (Fermentas) according to the manufacturer’s instructions and diluted 1:10 with RNase free water. qRTPCR was performed as previously described ^39^. Briefly, qRTPCR reactions were prepared with Quanta SYBR (Quanta Bio) and PCR was performed on a Step One Plus device (Applied Biosystems). Data was analyzed using ΔCt method, Step One Plus software package (Applied Biosystems) and Excel (Microsoft). Endogenous control was GAPDH. Primers were designed by qPrimer Depot. The details of the primer are listed in the table in supplementary file.

### Transcervical mouse infections and determination of bacterial burden

ASCT2/SLC1A5 KO mice generated in a C57BL/6 background were obtained from the Australian National University (ANU) ^25^. All animal experiments were performed in accordance with protocols approved by animal care and experimentation of German Animal Protection Law approved under the Animal (Scientific Procedures) Act 1986 (project license 55.2-2532-2-762). The mice used for experiment were between 10-14 weeks old. Five days before transcervical infection, mice were treated subcutaneously with 2.5□mg of DepoProvera (medroxy-progesterone acetate). The mice were transcervically infected with 1□×□10^7^ infection-forming units of *Chlamydia* using a non-surgical embryo transfer device (ParaTechs Corp.). The mice were euthanized 7□days post-infection and the uterine horns were taken for further analysis. The uterine horns were homogenized in SPG buffer and DNA was isolated using DNeasy blood and tissue kit (Qiagen). Quantitative PCR was used to enumerate *Chlamydia* and host genome copy number. The following primers were used for amplifying the *C. trachomatis lytA* gene that was cloned into the vector: fwd, 5′-TCTAAAGCGTCTGGTGAAAGCT-3′ and rev, 5′-GAAATAGCGTAGTAATAATACCCG-3′. Normalization of bacterial genome to that of the host was performed using mouse synectin primers: fwd, 5′-ACTAATGTCAAGGAGCTGTACG-3′ and rev, 5′CCTCCGACTTGAACACTTCC-3′. Quantitative PCR with reverse transcription (RT–PCR) was performed as described below. Data were analyzed using Step One Plus software package (Applied Biosystems) and expressed as the ratio of chlamydial genome to host genome (*lytA/*synectin). GraphPad Prism 7 was used to generate a scatter column chart and perform statistical analysis. One-way analysis of variance (ANOVA) with Newman– Keuls multiple-comparison tests was performed with the significance level set to less than 0.01. Statistical analysis was performed to decide the sample size used in mouse infection by the Institute of Mathematics, University of Würzburg under the allowance A2 55.5-2531.01-49/12. All mouse experiments were carried out with five female mice per treatment group. Mice in each experiment were age-matched and cage mates were randomly distributed into different treatment groups to avoid cage effects.

### Statistical analysis

In all experiments, a minimum of three technical replicates was used and the *n* number refers to the number of independent experiments performed. The data are presented as box plots with the mean and s.e.m. Statistical analyses were performed with the Prism 7.2 package (GraphPad Software). ExactTest () function as a part of edgeR module in R 3.3.4 was used to carry out pairwise comparison and calculate p-values. False discovery rates were calculated as q-values using Benjamini-Höchberg algorithm implemented in edgeR module.

## Author contribution

KR and TR designed the experiments. The experiments were performed by KR, NV and TW. RNA-Seq was performed and analyzed by KR, EW and AB. Samples for Mass Spectrometry were prepared by KR and SJ. NV, CH, MS and WE performed the metabolic flux analysis of axenic culture and data interpretation. SJ, WS and AS performed metabolic flux analysis in host cell culture. FD provided plasmids and cell lines. Click reagents were synthesized by JF and JS. KR and TR wrote the manuscript.

## Acknowledgment

We kindly acknowledge Stefan Broer and Anselm Enders, Australian National University for providing the SLC1A5 KO mice. We acknowledge Claudia Gehring and Daniela Bunsen for processing the TEM samples. We thank Vera Kozjak-Pavlovic for the cell line expressing shRNA against GLS1 and Andreas Demuth for critically reading the manuscript. KR was partially funded by the Frauenbüro in the frame of the “Qualification for junior scientist for professorship programme”. This work was also supported by the German Research Foundation (DFG) grant WO 2108/1-1 to WE and the GRK 2157 “3D-Infect” to TR. AB was supported by grants of the German Excellence Initiative to the Graduate School of Life Sciences, University of Würzburg. SJ was supported by the DFG grant SCHU2670/1-1.

## Supplementary data

**Table 1:**
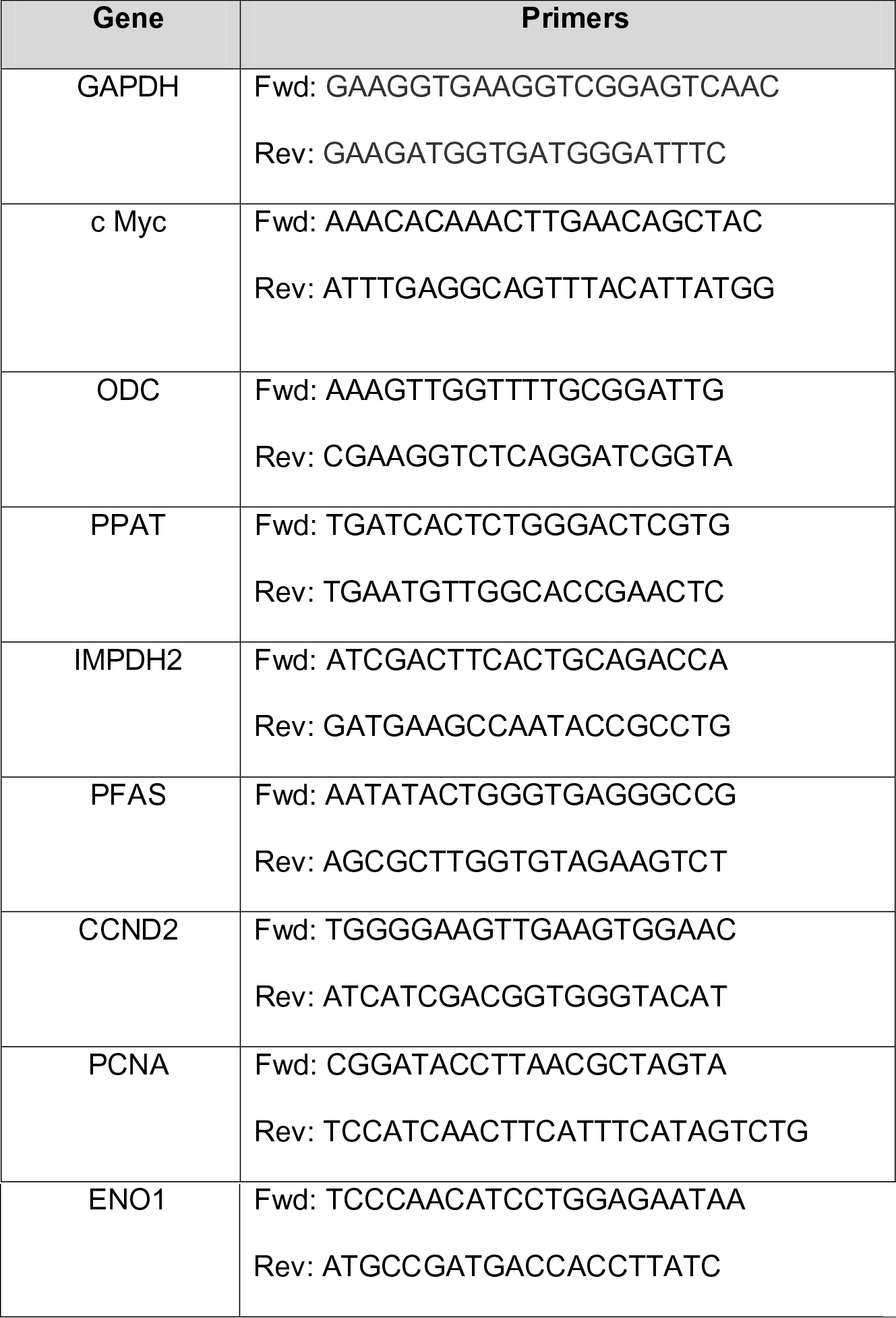
Primers for RT-PCR

## Supplementary Fig

**Fig. S1: Glutamine metabolism in *Chlamydia*. a.** Chlamydial EBs were purified and incubated in axenic media containing G6P with or without Gln. After 2 h NADH levels were measured as described in the Method section. * Indicates p value <0.05 and ** Indicates p value <0.01. n=3. **b.** Renografin-purified EBs were incubated in axenic medium with ^15^N-labeled glutamine specimens and analyzed via GC-MS (see Methods for details). **c.** Renografin purified EBs were incubated with either G6P or G6P and glutamate or glutamine for 2 h. The bacteria were then used to infect freshly plated HeLa cells. 24 hpi, the inclusion forming units were calculated and plotted in the graph. The number of inclusions in G6P and glutamine-treated sample was normalized against G6P-treated sample. *** Indicates p value <0.001, n=3. **d.** Scheme showing the shuttling of ^13^C and ^15^N flux from U-[^13^C_5_] Gln and ^15^N (amine) Gln from the axenic culture into Chlamydia for DAP biosynthesis.

**Fig. S2: Chlamydial infection triggers altered glutamine metabolism in infected cells.** Diagram outlining the differential use of glutamine (Gln) by the metabolism of host and chlamydial cells. Host cells feed Gln-derived α-ketoglutarate (α-KG) into their complete TCA cycle. This leads to the production of M+4, M+2 and M+1 isotopologues of all TCA cycle intermediates generated by the first, second and third round of the cycle, respectively. Alternatively, host cells can also use reductive carboxylation of Gln-derived α-KG to generate M+5 citrate (Cit), which is further converted to oxaloacetate (Oxa) by the host cell metabolism. The different isotopologues of Oxa are then used to generate M+4, M+3, M+2 and M+1 labelled aspartate (Asp). As no Gln-derived labelling of pyruvate was observed in infected cells, it can be concluded that the gluconeogenesis enzymes required for the conversion of oxaloacetate to pyruvate are inactive. In contrast, the truncated TCA cycle of *Chlamydia* can only produce M+4 isotopologues of α-KG, succinate (Suc), fumarate (Fum), malate (Mal) and Oxa, resulting in M+4 labelled Asp. Infected cells increase the uptake of Gln from the culture media and enhance its catabolism to provide metabolic intermediates for bacterial growth.

**Fig. S3: a.** HeLa229 were left uninfected or infected with *Chlamydia* at an MOI of 1 for 24 and 36 h. RNA was isolated and RT-PCR was performed to analyze the levels of c-Myc target genes (n=3). Fc indicates the fold change with respect to uninfected cells. **b/c/d.** Human fimb **(b)**, mouse fimb **(c)** and U2OS cells **(d)** were infected with *C. trachomatis* serovar L2 at an MOI of 1 for various time points. The cells were lysed and analyzed for c-Myc expression via Western blot. ß Actin serve as the loading control and cHSP60 indicates the infection level. n=3. **e/f/g.** HeLa229 cells were infected with either *C. trachomatis* serovar D **(e)**, *C. pneumoniae* **(f)** or *C. muridarum* **(g)** for different time points as indicated and the cell lysate were analyzed for c-Myc expression by Western blot. ß Actin serves as the loading control and cHSP60 indicates the infection level. n=3. **h.** HeLa229 cells were infected with *Chlamydia* at an MOI of 1 for different time points. Total RNA was isolated and levels of c-Myc mRNA were detected using RT-PCR. n=3, *** indicates p value <0.001. **i.** HUVEC cells were infected with *Chlamydia* for 24 hpi, the cells were fixed with PFA and immunostained for c-Myc (green), cHsp60 (red) and phalloidin (blue). All experiments were done in three technical replicates. **j.** Hela229 cells were either left uninfected or infected with *Chlamydia* at an MOI of 1 for 24 or 36 hpi. The cells were lysed and the cytoplasmic and nuclear fraction was isolated as explained in the Methods section. The lysate was analyzed using Western blotting for c-Myc. GAPDH was used as a control for cytoplasmic fraction and Histone for nuclear fraction.

**Fig. S4: a.** U2OS cells were treated with different concentrations of anhydrous tetracyclin (AHT) to investigate adverse effects of AHT on chlamydial infection. The cells were then infected with *Chlamydia* at an MOI 1 for 30 h. The cells were lysed and analyzed by Western blotting for cHsp60 (infection level) and ß Actin (loading control). n=3. **b.** Infectivity assay from the experiment shown in Fig. 5A and 5B. AHT-induced and/or U0126/Ly294002 cells were infected with *Chlamydia* for 30 h and the cells were lysed with glass beads. Different dilutions of the lysate were used to infect freshly plated HeLa229 cells. After 24 h the cells were lysed and the infection level (cHsp60) was detected by Western blotting. ß Actin serve as the loading control. **c.** The number of inclusions from the above experiment was counted to plot the graph. n=3. * Indicates p value <0.05 and ** Indicates p value <0.01. **d.** HeLa229 cells carrying an inducible shRNA expression cassette to silence c-Myc expression were either induced with AHT or left uninduced. The cells were infected with *Chlamydia* at an MOI 1 for 24 h. The cells were lysed and analyzed using Western blotting. cHsp60 indicated the infection level, ß Actin served as loading control. n=3.

**Fig. S5: a,b.** Gene seq enrichment analysis using the MSigDB C5 (GO Term) collection of genes sorted according to regulation between cells infected with *Chlamydia* for 12 h (a) or 12, 24, 36 h (b) and uninfected conditions. Enrichment plots and NES values of gene sets involved in amino acid transport and metabolism are shown as an example. NES: normalized enrichment score. **c.** Human Fimb cells were grown in basic formulation of DMEM containing 4.5 g/L D glucose and different concentrations of L-glutamine (Gln). These cells were then infected with *Chlamydia* at an MOI of 1 for 24 h. Chlamydial growth was examined by Western blotting. **d.** HeLa229 cells were grown in in basic formulation of DMEM containing 4.5g/L D glucose and incubated with different concentrations of glutamine and infected with *Chlamydia* at an MOI 1 for 30 h. The cells were lysed and analyzed by western blotting. cHsp60 indicates the infection level ß Actin serve as the loading control. n=3. **e** Infectivity assay were performed using cell lysates from Fig. 5a at different dilutions to infect fresh HeLa229 cells. 24 hpi the cells were lysed and analyzed by Western blotting for chlamydial Hsp60 and ß Actin. **f.** HeLa229 cells were either grown with or without glutamine/nucleosides as supplements. The cells were left uninfected or infected with *Chlamydia* at an MOI 1 for 30 h. The cells were lysed and analyzed by western blotting. cHsp60 indicates the infection level. ß Actin serve as the loading control. n=3.

**Fig. S6: a.** HUVECs were either left uninfected or infected with *Chlamydia* at an MOI 1 for 24 h. RNA was isolated to detect the levels of various transporters. n=3. **b.** Infectivity assay from Fig. 6i. HeLa229 cells carrying an inducible shRNA expression cassette to silence c-Myc expression were either induced with AHT or left uninduced. These cells were in addition transfected with an expression plasmid for SLC1A5. The cells were infected with *Chlamydia* with an MOI 1 for 30 h. The number of inclusions was counted to plot the graph or the cell lysate was used to analyze cHsp60 levels via Western blotting. **c.** Western blot analysis of SLC1A5 in BL6 and SLC1A5 KO mice. Tissue lysate from 5 mice used in the experiment were used for the analysis. ß Actin is used as the loading control.

