## Supplementary figures and images for "A central role of glutamine in *Chlamydia* infection"

### Supplemental Figure 1

##### Supplementary Fig. 1

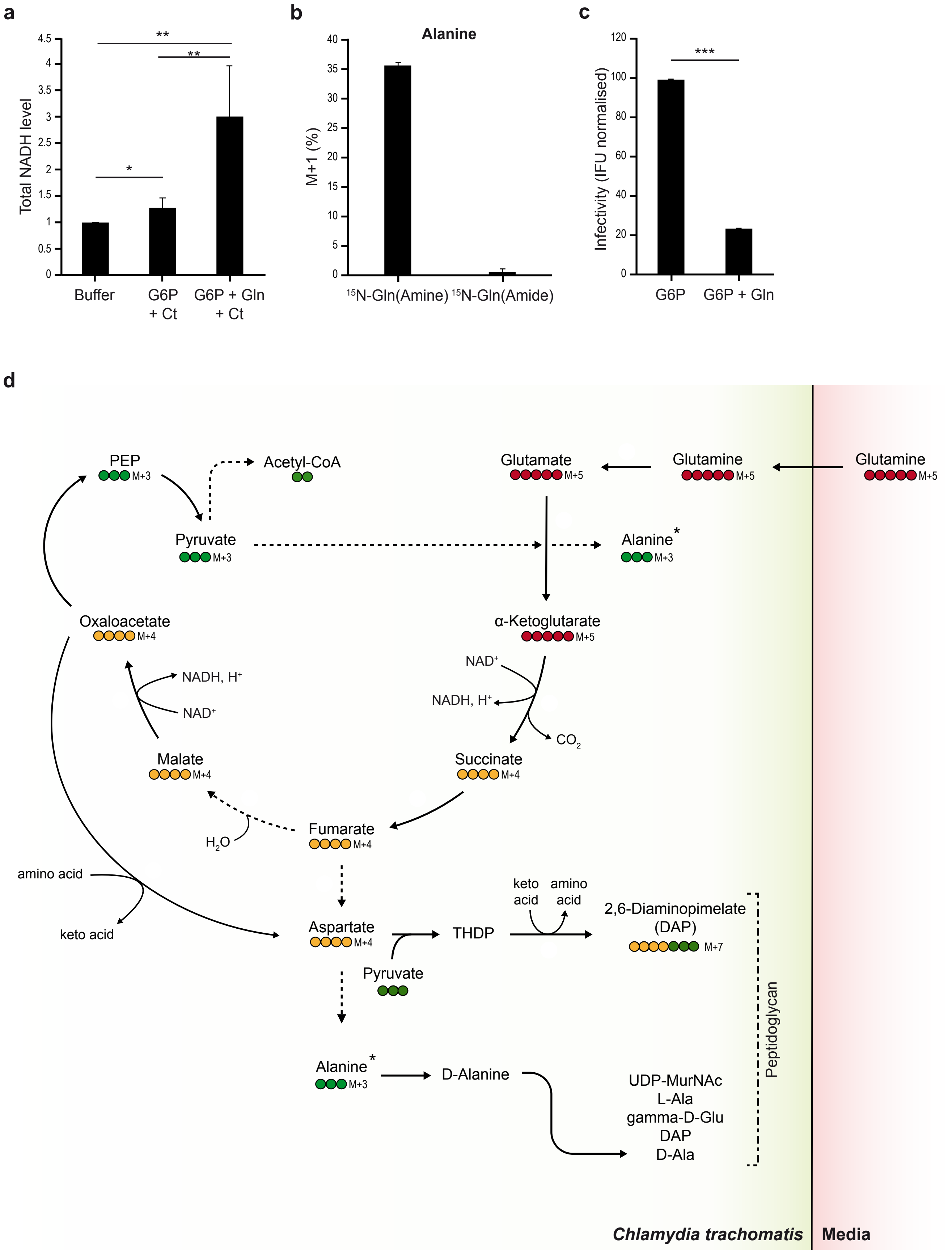

### Supplemental Figure 2

#### Supplementary Fig. 2

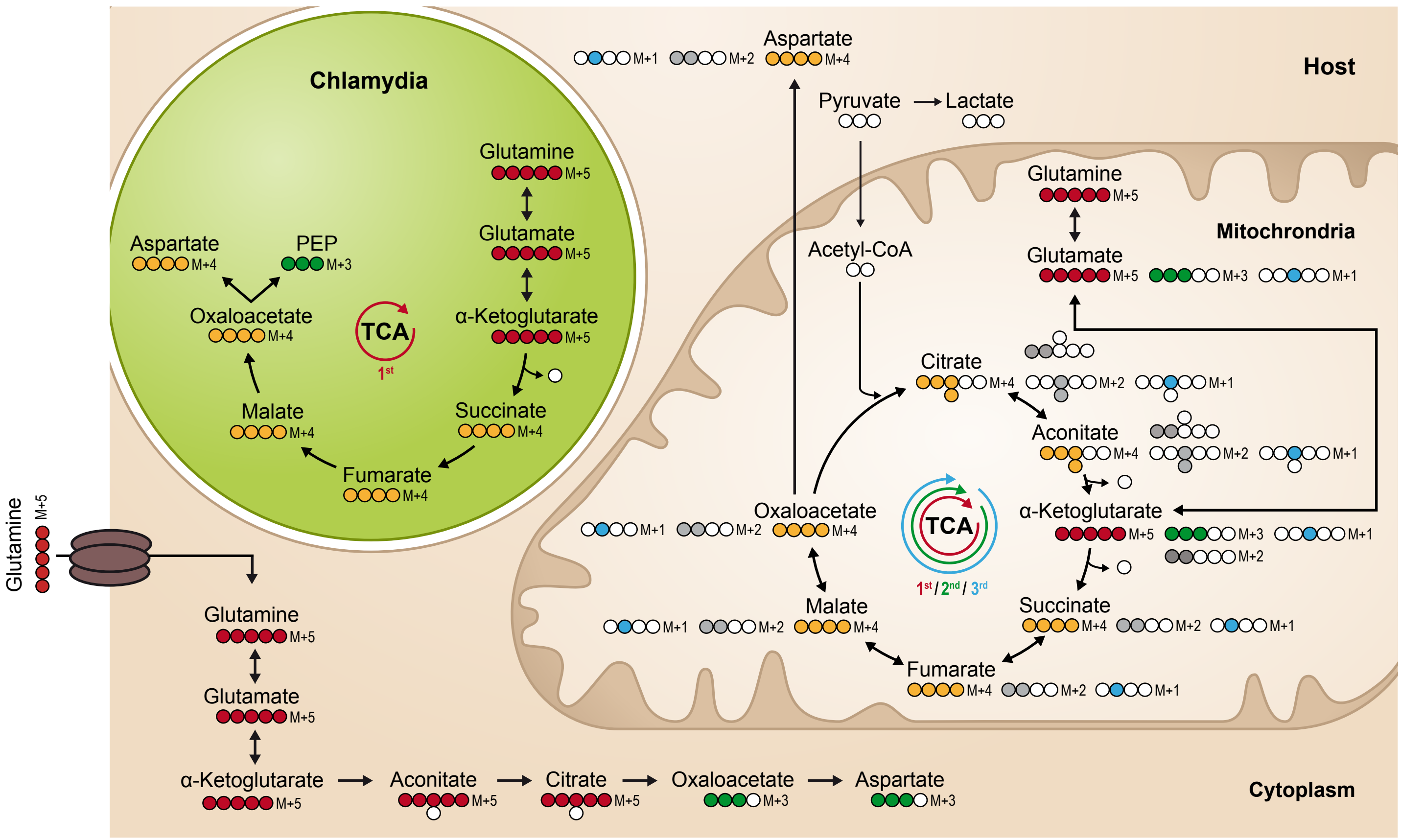

### Supplemental Figure 3

**Supplementary Fig. 3**

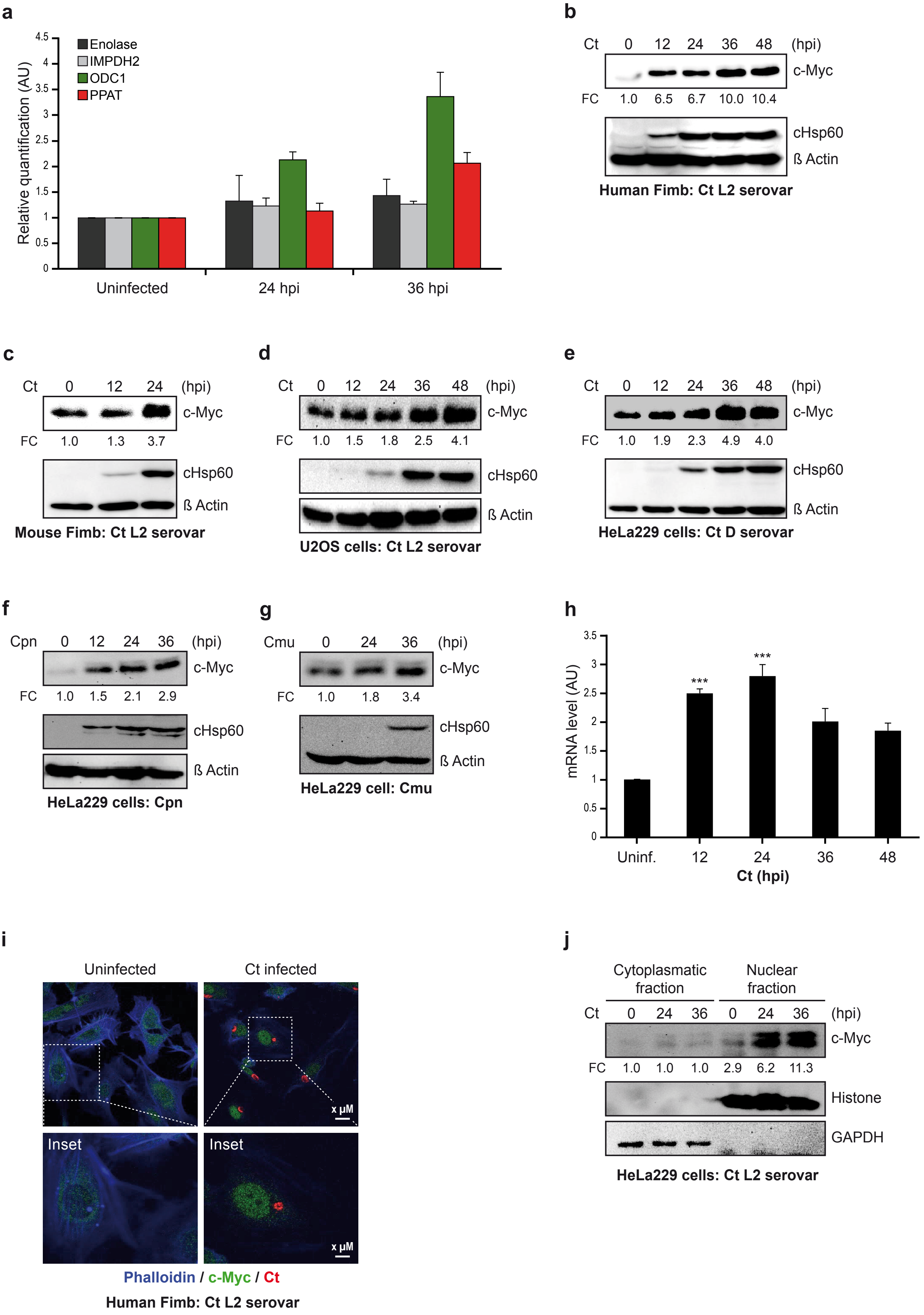

### Supplemental Figure 4

#### Supplementary Fig. 4

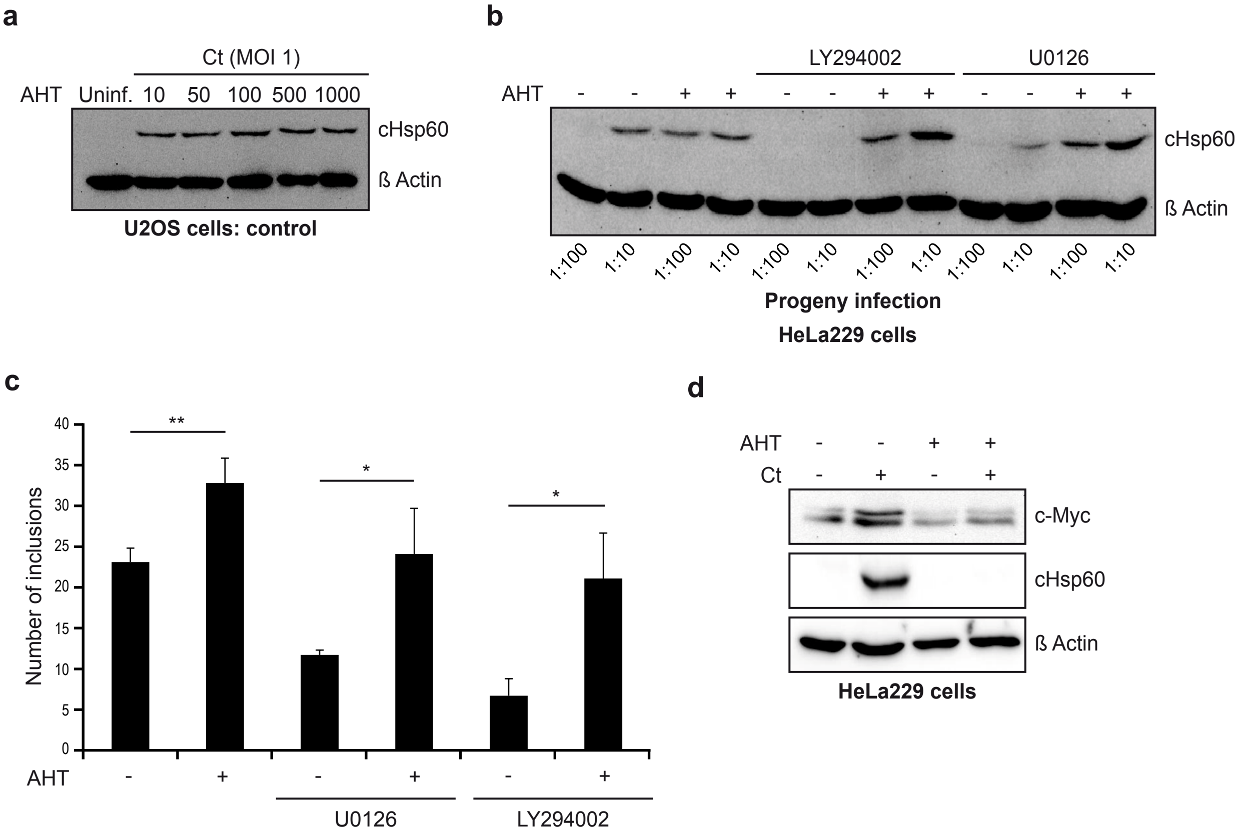

### Supplemental Figure 5

##### Supplementary Fig. 5

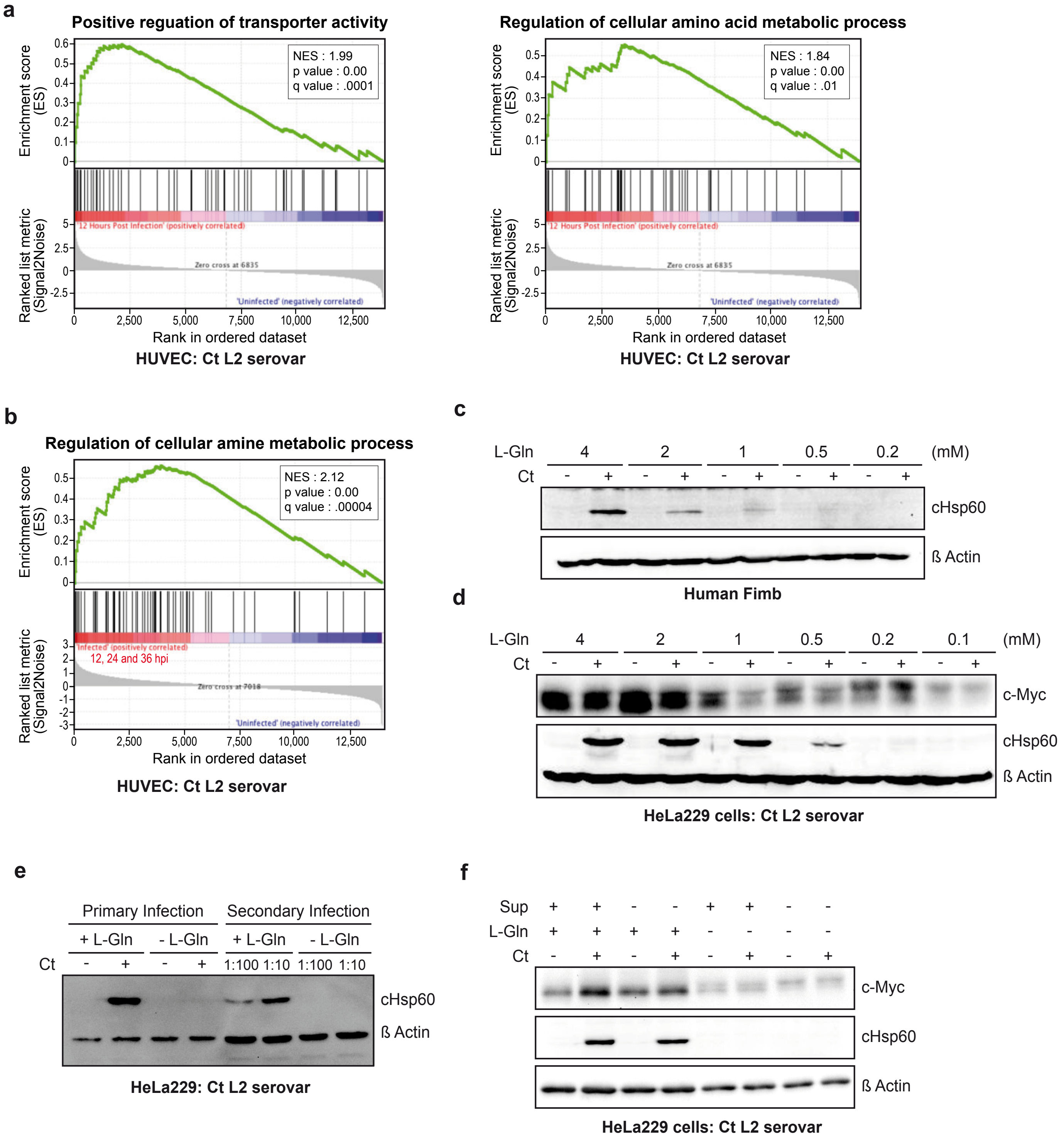

### Supplemental Figure 6

##### Supplementary Fig. 6

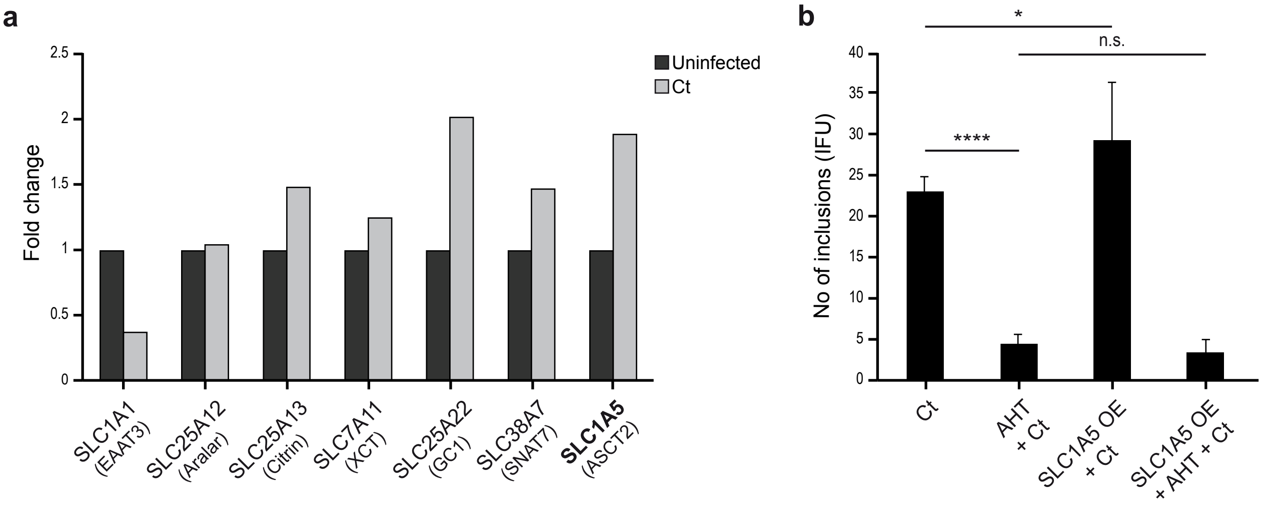


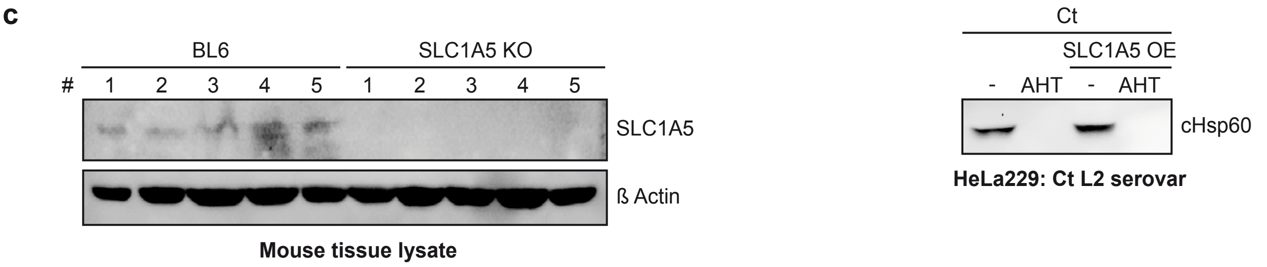
